# Low-Dimensional Frontal Feedback Resolves High-Dimensional Visual Ambiguity in Human Visual Cortex

**DOI:** 10.64898/2026.03.23.713175

**Authors:** Yiyuan Zhang, Jirui Liu, Jia Liu

## Abstract

One major distinction between artificial neural networks and biological brains is the prevalence of extensive, long-range feedback connections in biological systems. Here we investigate unique contributions of these hierarchical feedback signals beyond feedforward processing and local recurrence by exploring their mechanistic role in resolving visual ambiguity caused by occlusion. Both empirical fMRI and EEG experiments and computational modeling show that when sensory evidence for faces became insufficient, the ventrolateral prefrontal cortex (vlPFC) sustained a low-dimensional belief state (e.g., animate vs. inanimate objects) and transmitted this abstract information back to the animacy map in the ventral temporal cortex (VTC) encompassing face-selective representations. Critically, in the hierarchical vision model inspired by this finding, this frontal feedback did not reshape the attractor geometry of the VTC; instead, it provided guidance to reroute ongoing neural dynamics away from ambiguous pseudo-states toward face attractor basins in the energy representational landscape. This control-based mechanism of feedback signals thus enabled perceptual completion by reconstructing missing facial features with temporal costs verified through EEG. Together, this multimodal study bridges analysis-by-synthesis theories of vision and dynamical-systems perspectives on long-range feedback as state-space control, and offers inspiration for the design of hierarchical AI architectures incorporating feedforward, recurrent, and feedback connections.

## Introduction

The primate visual system is traditionally viewed as hierarchically organized, in which nearly every ascending feedforward pathway is complemented by extensive feedback projections ^1,2^, especially long-range connections originating from frontal cortex and projecting back into cortical regions of the ventral visual stream (Figure 1A). Such extensive long-range feedback contrasts with contemporary artificial vision systems, where many state-of-the-art models rely predominantly on rapid feedforward computations, supplemented at most by shallow local recurrent processing ^3–6^. Given the substantial wiring length, metabolic cost, and processing delay associated with this hierarchical architecture, the pervasive presence of feedback pathways in the primate visual system raises a critical question: what unique computational function does long-range feedback serve in the primate brain?

**Figure 1.**
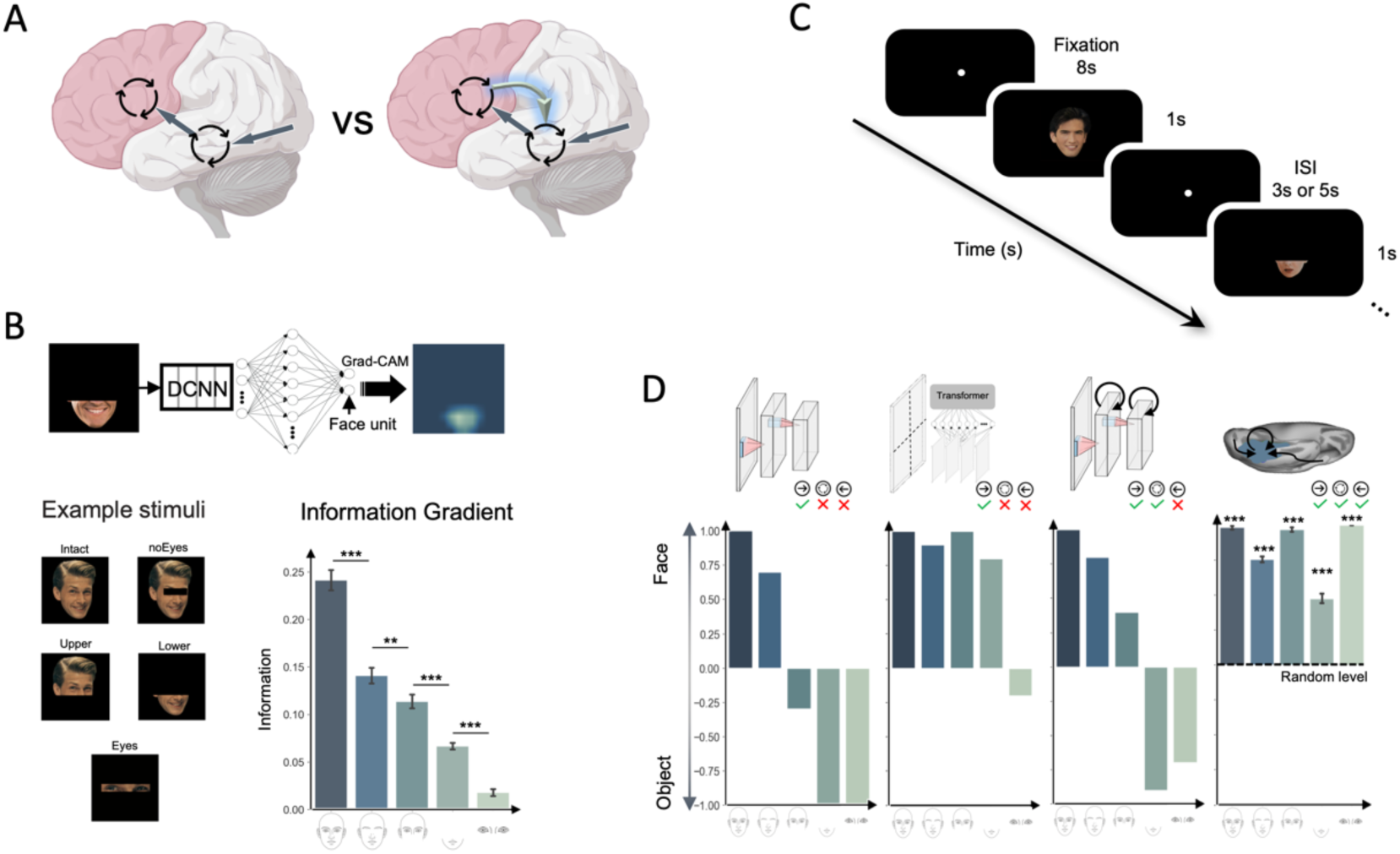
Long-range feedback signals are essential for recognizing occluded faces. (A) Schematic illustration of various connection types in the primate visual system. The right panel highlights extensive long-range feedback connections, in contrast to the predominantly feedforward connections with shallow local recurrence depicted in the left panel. Schematic illustrations were created by FigDraw (www.figdraw.com). (B) Methods for evaluating facial information content in stimuli and example faces from IGOF dataset. Top: A DCNN classifier was trained to distinguish faces from objects, and Grad-CAM was applied to quantify facial information. Bottom left shows example stimuli, including Intact Face (Intact), Eyes-Occluded Face (noEyes), Upper-Half Face (Upper), Lower-Half Face (Lower), and Eyes-Only Face (Eyes); Bottom right illustrates a systematic reduction in facial information indexed by heatmaps with increasing occlusion severity. Error bars denote standard deviation (S.D.). (C) Stimuli and experimental procedure of human fMRI experiment. ISI: interstimulus interval. (D) Decoding accuracy comparison of AlexNet (purely feedforward), ViT (state-of-the-art feedforward model with Transformer architecture), CORnet-S (feedforward + locally recurrent), and human FFA (presumably feedforward + locally recurrent + long-range feedback) under varying occlusion levels. Decoding accuracy was normalized between –1 (confident misclassification as non-face objects) and 1 (confident classification as faces). **: *p* < 0.01; ***: *p* < 0.001

A common assumption is that feedback helps stabilize perception under conditions of incomplete or degraded sensory inputs ^7–9^. When an object is partially hidden, the visible fragments may support multiple plausible interpretations, rendering recognition inherently underdetermined ^10,11^. This scenario highlights a critical divergence between biological and artificial vision: classical theoretical vision models ^1,12–16^ and deep convolutional neural networks (DCNNs) ^17–19^ perform remarkably well with intact stimuli yet exhibit pronounced vulnerability to occlusion ^20,21^, whereas primate vision remains comparatively robust ^20,21^. Converging behavioral and neurophysiological evidence suggests that this resilience involves additional processing time ^22,23^ and recruitment of cortical regions beyond ventral visual stream ^24–29^. The ventrolateral prefrontal cortex (vlPFC) is of particular interest because of its bidirectional connections with high-level visual areas ^30,31^ and its neurons selectively enhancing representations of occluded shapes in visual cortex ^24,25,29,32,33^. Within the predictive coding framework, such top-down projections are hypothesized to convey predictions that are iteratively compared with and corrected by bottom-up signals, yielding stable perceptual interpretations over time ^26,33–36^. Despite these advances, however, the mechanism of feedback projections remains poorly understood, particularly regarding the precise content carried by these long-range feedback signals and the dynamic interactions with the representation geometry of targeted visual cortex to reroute the system’s neural trajectories away from sensory ambiguity into a stable percept.

To address this question, we employed an integrative, multimodal approach that holds the visual evidence under control while giving us multiple perspectives into the same underlying computation. Faces are an especially informative test case because the ventral stream contains well-characterized face-selective representations ^37–39^, yet everyday occluders often remove precisely the features that are most diagnostic. Specifically, we first developed a set of information-graded occluded faces (Figure 1B) in which the available facial evidence is reduced in a systematic and parametric way, aiming to manipulate how much diagnostically useful information remains for categorization. Using functional magnetic resonance imaging (fMRI), we then examined interactions between frontal and temporal visual cortical regions ^24,29^, specifically focused on how functional connectivity between the vlPFC and ventral temporal cortex (VTC) varied as sensory evidence became increasingly insufficient, to identify the content carried by the feedback signals. We next built a biologically plausible hierarchical vision model consisting of a high-level vlPFC-like module and a low-level VTC-like module connected by their dynamic interactions to examine how feedback signals could resolve sensory ambiguities. Finally, we conducted a complementary electroencephalography (EEG) experiment to characterize the temporal dynamics of VTC activity during ambiguity resolution. Together, this multimodal approach aimed to establish a comprehensive understanding of the role of long-range feedback signals in stabilizing perception under occlusion.

## Results

### Frontal Feedback is Necessary for Recognizing Occluded Faces

When critical parts of an object are occluded, bottom-up sensory input alone becomes insufficient to reach a stable neural state, necessitating additional top-down feedback signals to stabilize perception ^24–26,29^ (Figure 1A). To systematically characterize the functional role of these feedback signals, we constructed a novel dataset of the Information-Graded Occluded Faces (IGOF, Figure 1B), which parametrically varied the extent of visible facial information. Previous occluded recognition studies typically focus on occlusion size as a measure of visual information loss ^20,25,40–42^, but the amount of usable information in faces varies across different facial features, with regions like the eyes and nose being crucial for face recognition ^43–46^. To objectively quantify the facial information retained in each image, we used a DCNN to identify features most critical for face recognition by training a classifier on the DCNN’s feature representations to discriminate faces from non-face objects (see Methods for details). Given the representational similarity between DCNNs and the human feedforward visual pathway ^15,16,47–50^, the critical facial features identified by the DCNN provide a biologically plausible foundation for designing controlled information gradients. Subsequently, using the Gradient-weighted Class Activation Mapping (Grad-CAM) method ^51^, we generated heatmaps highlighting the image regions essential for face categorization (Supplementary Figure 1). The average heatmap intensity within facial regions thus provided an objective measure of facial information available in each stimulus, enabling a parametric assessment of the occlusion level at which the VTC transitions from stable, category-specific representations to ambiguous states.

Following this approach, we constructed five occlusion conditions (Figure 1B, examples at bottom left) with progressively reduced facial information, ranging from fully visible faces (Intact condition) to images with critical facial components commonly occluded in real-world scenarios. These conditions included faces with occluded eyes like wearing sunglasses (noEyes condition), faces with only the upper half visible like wearing a mask (Upper condition), faces with only the lower half visible as if the upper face is covered by a hat or shadow (Lower condition), and faces showing exclusively the eye region like wearing a veil (Eyes condition). Using the DCNN-derived metric, we confirmed a systematic reduction in facial information for face recognition across successive occlusion levels (one-way ANOVA *F*(4,95) = 139.65, *p* < .001; Tukey HSD *p*s < .01; Figure 1B, bottom right), verifying that the IGOF dataset effectively provided a controlled gradient of facial information from intact to severely impoverished visual stimuli.

With this stimulus set, we evaluated how the human brain and artificial vision models performed in recognizing partially occluded faces. Figure 1C outlines the human fMRI experimental procedure, which used a rapid event-related design, where 30 human participants viewed images from the IGOF dataset alongside everyday tools, while brain activity in the fusiform face area (FFA ^37^) was recorded. Figure 1D compares the performance of three artificial neural networks with that of human FFA decoding accuracy. Performance was quantified using a normalized decoding accuracy ranging from –1 to 1, where 1 indicated confident classification as a face, –1 indicated confident misclassification as a non-face object, and 0 represented chance-level performance. The left panel of Figure 1D shows results from AlexNet ^52^, a classic feedforward DCNN without recurrent or feedback connections. AlexNet’s face decoding accuracy substantially declined with increased occlusion (Figure 1D, left). Notably, AlexNet classified severely-occluded faces (Lower and Eyes conditions) as non-face objects, underscoring its fragile performance under substantial sensory ambiguity. This kind of vulnerability also exists in other classic feedforward neural networks (Supplementary Figure 2). In addition, we tested a state-of-the-art feedforward model, Transformer-based ViT that is capable of maintaining about 60% top-1 accuracy on ImageNet images with ∼80% of the pixels randomly occluded ^53^. The ViT also failed in the most severe occlusion (Figure 1D, second to the left). The third panel shows the performance of CORnet-S ^3^, a neural network integrating shallow recurrent connections to enable iterative processing. CORnet-S demonstrated slightly improved robustness compared to AlexNet in moderately occluded faces (Upper condition) but still showed substantial impairments under severe occlusions (Lower and Eyes conditions). In contrast, the right panel of Figure 1D and Supplementary Figure 3 show the resilience of the FFA’s decoding performance across all occlusion levels. Multivariate pattern analysis of the fMRI data revealed that the FFA reliably distinguished faces from non-face objects across occluded conditions (*p*s < .001, Bonferroni corrected). In addition, the classic GLM showed that the FFA had significantly higher beta values for various occluded faces than those for non-face objects (Supplementary Figure 4). This robust representational stability contrasts with performance degradation observed in both purely feedforward and shallow recurrent neural networks.

Together, these results indicate that the human FFA maintains robust face representations despite substantial loss of bottom-up sensory information, suggesting reliance on prior experience or context provided by top-down feedback from higher cortical areas. Next, we investigated the cortical origin of this predictive feedback signal and the mechanism supporting robust perception under conditions of insufficient feedforward input.

### Low-Dimensional Feedback from vlPFC Stabilizes VTC Representations

Neurophysiological studies on nonhuman primates implicate the vlPFC in providing top-down feedback signals during visual recognition under conditions of occlusion ^24,25^. Accordingly, we hypothesized that human vlPFC similarly generates feedback signals to support visual cortex when processing degraded sensory input. To test this hypothesis, we measured functional connectivity between the vlPFC and FFA as facial occlusion progressively increased. Surface-based searchlight analyses revealed that as occlusion severity intensified there was a significant increase in both the proportion of cortical vertices in the vlPFC functionally coupled with the FFA (*R²* = 0.67, *p* < .001) (Figure 2A and Figure 2B left) and the strength of vlPFC–FFA connectivity (*R²* = 0.19, *p* < .001) (Figure 2B right). In other words, vlPFC engagement was heightened in response to increasing occlusion, suggesting that frontal cortex dynamically modulates its interactions with visual areas when sensory evidence becomes insufficient. This finding contrasts with traditional feedforward models of visual hierarchy; instead, it aligns with an interactive, predictive framework, in which higher-order cortical regions proactively provide information to facilitate visual recognition ^34,54^.

**Figure 2.**
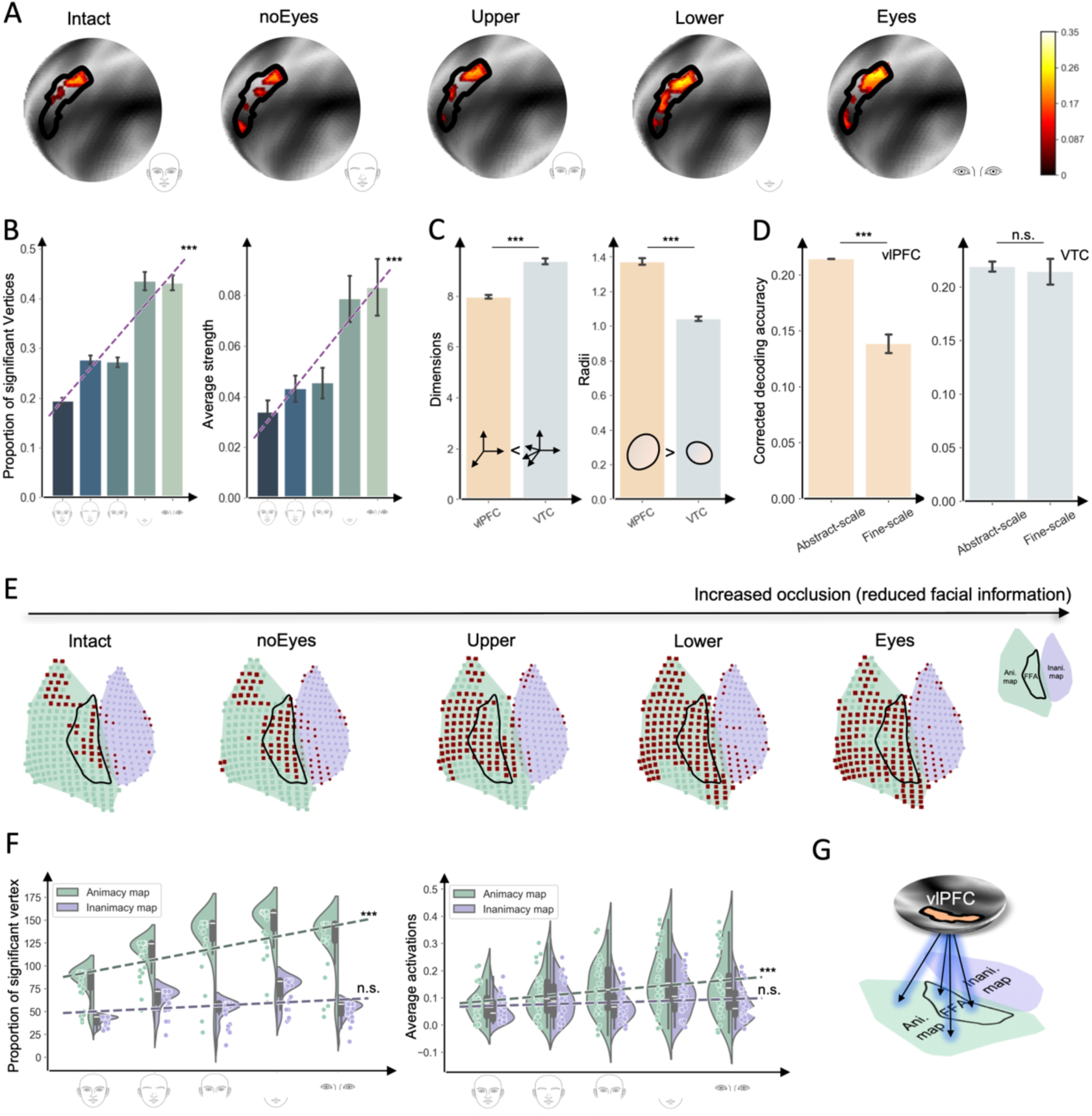
Feedback from the vlPFC modulated the animacy map in the VTC through functional connectivity. (A) Cortical vertices in the vlPFC showing significant functional connectivity to the FFA across occlusion severity. Black contour outlines the vlPFC in the HCP-MMP template ^55^; Dark and light grays indicate sulci and gyri, respectively; Color bar indicates connectivity strength. (B) Proportion of cortical vertices in the vlPFC functionally connected to the FFA (left) and their connectivity strength (right) across occlusion severity. Linear regression analysis revealed significant correlations, indicating that vlPFC engagement was heightened in response to increasing occlusion. (C) Effective dimensionality (left) and representation radius (right) of representational manifolds in the vlPFC and VTC, respectively. Notably, the lower dimensionality and larger representation radius of vlPFC manifold indicate its representation is more abstract, encoding a broader range of stimulus categories than VTC representation. (D) Multivariate decoding analysis reveals the vlPFC encodes abstract semantic information (animacy vs. inanimacy) better than individual object categories (e.g., faces, chairs) (left panel), whereas the VTC showed no preference (right panel). Error bars indicate standard error of the mean (SEM). (E) Cortical vertices in the VTC (from the HCP-MMP template) showing significant functional connectivity with the vlPFC across occlusion levels. Each dot denotes a cortical vertex, with significant connectivity strength indicated by red (Bonferroni corrected *p* < 0.05 and correlation values are greater than 0.15). The animacy and inanimacy maps are colored in light green and purple, respectively, with the FFA outlined in black. Notably, vlPFC feedback predominantly targeted the animacy map, rather than specifically the FFA or diffusely influencing the inanimacy map. (F) Left: the proportion of functionally connected vertices in the animacy map increased significantly with decreasing facial information, while not in the inanimacy map. Right: significant increases in connectivity strength to the vlPFC as a function of increased occlusion were observed only in the animacy map, not in the inanimacy map. (G) Schematic illustration of reciprocal vlPFC-VTC interactions. Insufficient facial information from the FFA activated the vlPFC, whereas vlPFC feedback conveying abstract information of animacy activated the animacy map in the VTC, biasing perception toward the superordinate object category to which face category belongs. ***: *p* < 0.001

Having established the role of human vlPFC in feedback modulation, we investigated the nature of the information conveyed through these projections. To do this, we characterized neural manifolds constructed from population activity of vertices in the vlPFC and the VTC, respectively, to assess the geometry of neural representations. One key metric for manifolds is effective dimensionality ^56^, which indicates the level of abstraction in the representation, with lower dimensionality corresponding to higher abstractness ^57^. We found that vlPFC representations had significantly lower effective dimensionality than VTC representations (vlPFC: 7.96 dimensions; VTC: 9.36 dimensions; *t*(29) = –11.03, *p* < .001; Figure 2C left), suggesting that the vlPFC compresses visual information into fewer, more abstract dimensions. Additionally, the representational radius, which reflects the overall span of representations, was significantly larger in the vlPFC than in the VTC (vlPFC: 1.37; VTC: 1.04, arbitrary units; *t*(29) = 15.00, *p* < .001; Figure 2C right), suggesting that vlPFC representations are more abstract, encoding a broader range of object categories than VTC representations ^58–60^.

Given that vlPFC representations were more abstract than those in the VTC, and the fact that the FFA is part of the animacy map, which preferentially responds to animate stimuli (e.g., animals, faces, bodies) ^61–63^, we hypothesized that the vlPFC likely encodes a more abstract category of animate objects in general, rather than fine-grained objects (e.g., faces) in particular, compared to the VTC. To test this hypothesis, we conducted multivariate decoding analyses using another object-vision dataset ^64^, comparing the ability of the vlPFC and VTC to decode object categories at both abstract (animate: faces and cats vs. inanimate: houses, bottles, scissors, shoes, and chairs) and individual category levels (i.e., 7 categories). We found a significant two-way interaction of region (vlPFC vs. VTC) by level (abstract vs. individual levels) for decoding accuracy (two-way ANOVA *F*(1, 76) = 26.37, *p* < .001), indicating that differences in decoding accuracy between abstract and individual levels varied across brain regions. Furthermore, within the vlPFC, we found that vlPFC activity enabled significantly better decoding of abstract categories (animate versus inanimate, 21.43%, corrected for baseline) than individual categories (13.45%, corrected for baseline; mean difference = 7.98%, *p* < .001, Tukey HSD; Figure 2D left). In contrast, VTC decoding accuracy showed no preference for abstract categories (abstract vs. individual categories: 21.79% versus 21.87%, corrected for baseline; Figure 2D right). These findings suggest that the vlPFC preferentially supported abstract representations, while the VTC performed equally well across all levels.

Another way to examine the information conveyed by vlPFC feedback is to investigate which specific cortical regions the feedback targets within the VTC. We found that vlPFC feedback selectively modulated the animacy map under conditions of severe occlusion, rather than targeting task-specific FFA or uniformly affecting the whole VTC. That is, as facial information decreased, the vlPFC preferentially strengthened its functional connectivity with the animacy map, while connectivity with the inanimacy map remained relatively unchanged (Figure 2E). Quantitative analyses confirmed that both the proportion of functionally connected vertices within the animacy map (*F*(1,28) = 10.37, *p* < .05, *R²* = 0.69; Figure 2F left) and their mean connectivity strength (*F*(1,28) = 11.29, *p* < .05, *R²* = 0.79; Figure 2F right) significantly increased with decreasing facial information. In contrast, there was no significant change in either the proportion of connected vertices between the vlPFC and the inanimacy map (*F*(1,28) = 0.54, *p* = 0.52, *R²* = 0.15; Figure 2F left), or the strength of the connections (*F*(1,28) = 0.87, *p* = 0.42, *R²* = 0.23; Figure 2F right). Similar results were found in the other hemisphere of the brain as well (Supplementary Figure 5).

Therefore, when bottom-up facial information was insufficient, the vlPFC selectively enhanced the animacy map via low-dimensional feedback, conveying abstract information about animate objects rather than face-specific details (Figure 2G). This feedback thus likely biased perception toward superordinate object categories to which individual categories (i.e., faces) belong, thereby resolving ambiguity ^34,54,65^. Specifically, these fMRI results provide two biological constraints on any mechanistic account: (i) feedback signals should be low-dimensional and abstract (superordinate levels), and (ii) should preferentially couple to large-scale topography in VTC rather than fine-grained category maps. To further illustrate how such abstract feedback signals causally stabilize face representations under occlusion, we developed a biologically-inspired hierarchical computational model constrained by the above two principles that explicitly simulates the interactions between the vlPFC and VTC.

### Hierarchical Vision Model Illustrates the Generative Role of Frontal Feedback

The biologically-inspired hierarchical model that simulates the reciprocal interactions between the vlPFC and VTC was designed to formalize abstract feedback as a low-dimensional prior and to show how this feedback acted as a control signal that biases attractor dynamics under ambiguity. This model contains the VTC and vlPFC modules, which are interlinked through bidirectional connections (Figure 3A). Specifically, a pre-trained DCNN delivers bottom-up visual inputs to the VTC module, which is a recurrent neural network and forms face and object regions similar to human VTC (see Methods for details) ^57^. This initial step mimics early visual processing, where raw sensory data is converted into high-level feature representations. Subsequently, the VTC module sends feedforward signals to the vlPFC module, which is also a recurrent neural network, trained to extract abstract features as binary distinctions between animate and inanimate categories from the VTC input. Crucially, the vlPFC module does not merely reflect visual input but transforms this information into low-dimensional abstract representations, which are then fed back to the VTC module via top–down connections. This hierarchical architecture mirrors the division of labor observed in the human brain, where the vlPFC encodes superordinate categorical information (e.g. animacy vs. inanimacy) while the VTC processes detailed visual features, with iterative feedback loops enabling the inference of missing information.

**Figure 3.**
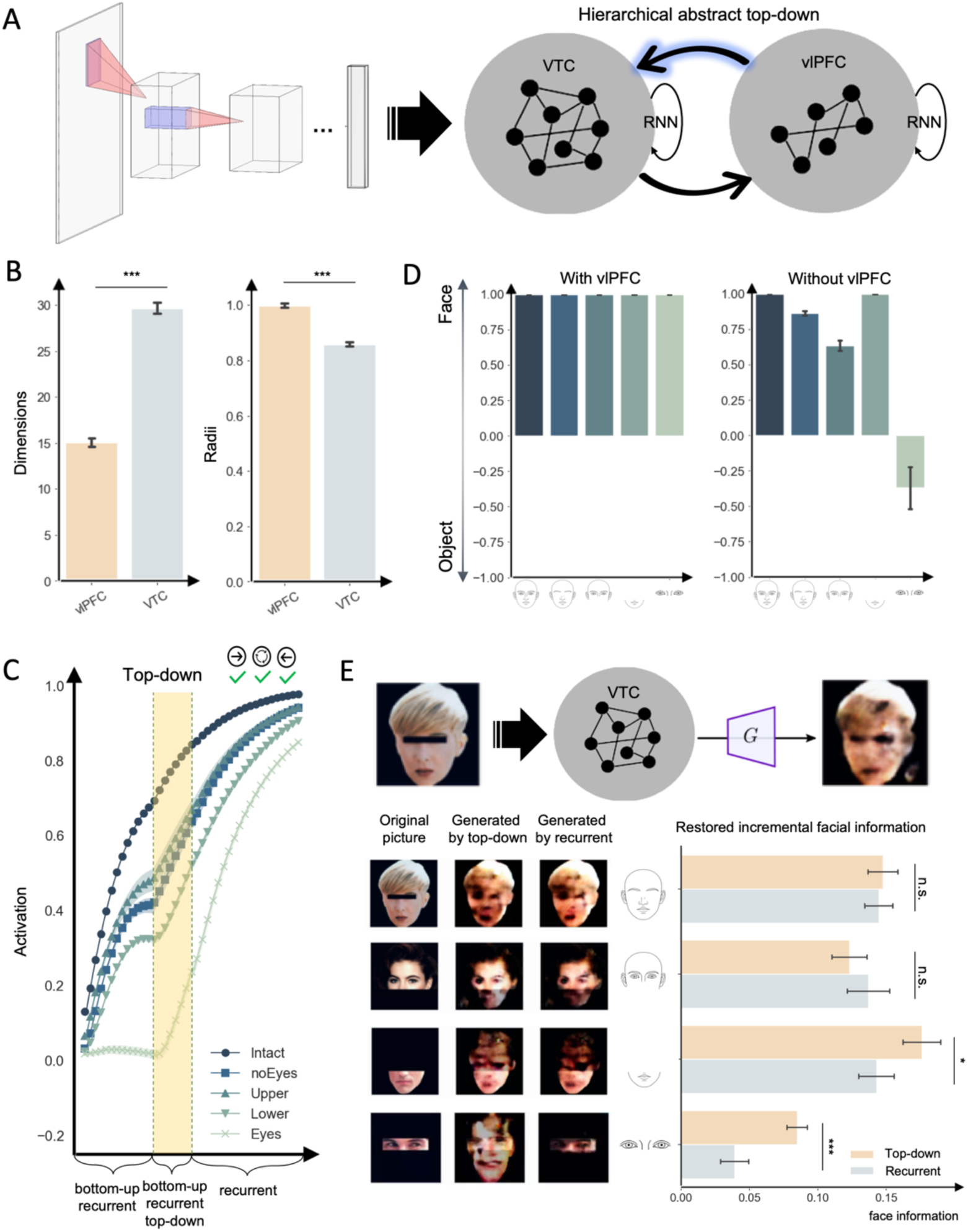
Feedback from vlPFC module helps generate missing facial information for occluded faces. (A) A hierarchical model with VTC and vlPFC modules. VTC module receives bottom-up input from DCNN, whereas vlPFC module receives bottom-up information from VTC module and then regulates VTC module through feedback connections. Both VTC and vlPFC modules are recurrent neural networks. (B) Effective dimensionality (left) and representation radius (right) of geometric manifolds for vlPFC and VTC modules, respectively, mirroring the geometric manifolds in human vlPFC and VTC. (C) Dynamics of face-specific activation (mean activation in the face region minus mean activation in the object region) in the VTC module with vlPFC feedback in each occlusion condition. Shaded areas indicate ± SEM. (D) Decoding performance of VTC module with or without feedback from vlPFC module. Left: with vlPFC feedback, the decoding accuracies for all occlusion conditions reached near-perfect performance. Right: without vlPFC module, less severely occluded faces were correctly but not perfectly decoded; for the most severely occluded faces, VTC module wrongly decoded it. (E) Generative decoding of representations in VTC modules. Top: Schematic of conditional GAN (cGAN) operation. After training, cGAN receives neural responses to occluded faces from VTC module and generate facial images. Bottom left: examples of original input images under each occlusion condition. With vlPFC feedback, most occluded facial regions are restored, generating a coherent face. Without vlPFC feedback and even with local recurrent connections, the quality of generated facial images deteriorates as occlusion increases. Bottom right: amount of facial information recovered by VTC module’s decoded images with/without feedback from vlPFC module at each occlusion level. *: *p* < 0.05; ***: *p* < 0.001

As expected, simulations showed that the model captured key differences in geometric manifolds between vlPFC and VTC modules, consistent with the observation in human brain (Figure 3B). Compared to VTC module, the geometric manifold of vlPFC module was significantly lower-dimensional (vlPFC module: 15.08 ± 0.41; VTC module: 29.70 ± 0.55; *t*(38) = 20.75, *p* < .001) and broader in radius (vlPFC module: 0.99 ± 0.01; VTC module: 0.86 ± 0.01; *t*(38) = 14.53, *p* < .001), indicating that vlPFC module maintains a low-dimensional abstract code while VTC module retains detailed, high-dimensional information. Accordingly, this model robustly recognized faces across all occlusion levels (Figure 3C and D left), which achieved near-perfect classification accuracy, even under the most severe occlusion conditions. This result replicates the robustness of human FFA decoding for faces with severe occlusion (Figure 1D right). In contrast, an ablated model without vlPFC feedback, which relied solely on bottom-up input and local recurrent processing, failed to reliably distinguish occluded faces from tools when critical facial information was missing (Figure 3D right; time series in Supplementary Figure 7A). Note that feedback from vlPFC module must be abstract, as when we trained vlPFC module to learn fine-scale category information rather than abstract representations, the decoding accuracies for occluded faces significantly decreased (Supplementary Figure 7B), similar to the decoding performance of the ablated model without feedback.

The necessity of feedback from vlPFC module is also illustrated by activation dynamics evolving over simulation time under different occlusion conditions (Figure 3C). In the ablated model, face-specific activation monotonically increases for faces in less severe occlusion conditions (e.g., Intact and noEyes conditions) but becomes low and unstable as occlusion severity increased. For the most severely occluded faces (Eyes condition), the ablated model fails to categorize them as faces (Figure 3D right and Supplementary Figure 7A). In contrast, with feedback from vlPFC module, decoding accuracies for faces with eyes only (Eyes condition) rapidly increase when feedback from vlPFC module is applied, and remains increased even after the feedback is removed. Thus, these activation dynamics illustrate that feedback from vlPFC module effectively restores and sustains face perception despite poor bottom-up sensory inputs.

Furthermore, the functional role of vlPFC feedback is generative in a dynamical sense, as it does not simply amplify existing activity, but actively redirects neural trajectories toward plausible solutions ^10,66^. When VTC module’s activity is decoded back into image space using a conditional generative adversarial network (cGAN) ^67^ (Figure 3E top), our model successfully reconstructs occluded face parts, progressively generating a coherent face despite large portions being missing (Figure 3E bottom left, middle column). In contrast, the ablated model’s decoding remains incomplete when occlusions are severe (Figure 3E bottom left, right column), emphasizing that vlPFC feedback does not merely amplify sparse bottom-up signals but to actively generate plausible missing features ^68,69^. To quantify this generative gain, we calculated the information recovered by models with and without vlPFC module under each condition. We found that the model with vlPFC feedback recovered significantly more information when decoding images via cGAN under severe occlusion compared to models without vlPFC module (Lower, *t*(38) = 2.25, *p* < .05; Eyes, *t*(38) = 5.22, *p* < .001) (Figure 3E bottom right). Furthermore, direct comparisons between generated images and their original counterparts reveal the same trend observed in the information content analysis: under severe occlusion conditions, images generated by the model with vlPFC module exhibit higher similarity to their original sources (Supplementary Figure 8). This finding illustrates the generative role of top-down feedback in perception, where vlPFC module incorporates high-level knowledge to supplement missing sensory details. To further examine whether abstract feedback influences the recurrent circuit of VTC module by reshaping representational basins, destabilizes competing solutions, or modulating trajectories through a fixed attractor landscape, we next analyzed the model’s state dynamics in an energy-representation space.

To reveal the mechanism by which abstract feedback from vlPFC module enables generative completion, we analyzed VTC state evolution in an energy-representation landscape ^57^. In this three-dimensional energy-representation manifold, X-Y plane embeds high-dimensional neural states into a two-dimensional representation, and Z-axis represents the energy of each state (Figure 4A left and middle). During network dynamics, neural states move toward low-energy regions (i.e., attractor basins) to form a stable perception. In our model, VTC module developed three major attractor basins (Figure 4A right): a face basin, a tool basin, and a pseudo-state basin that reflects visual ambiguity. Without vlPFC feedback, neural states of severely-occluded faces (e.g., Eyes condition) frequently fall into this pseudo-state basin, failing to converge to a clear category representation (Figure 4C; Figure 4E, a trajectory example of Eyes stimuli). With vlPFC feedback, the energy landscape remains unchanged as it is solely determined by VTC module’s recurrent connections. Instead, feedback alters the state trajectory within this landscape by providing additional driving force biased toward the animate superordinate category, thereby allowing the trajectory that would otherwise approach the pseudo-state basin to acquire sufficient energy to bypass it during state evolution (Figure 4B; Figure 4D, a trajectory example of Eyes stimuli; for all trajectories, see Supplementary Figure 9). This dynamic rerouting allows the trajectory to avoid settling into the ambiguous basin and instead converge along an appropriate path toward the face attractor basin. In this way, feedback from vlPFC module modifies the dynamics of VTC module to support a functional transition from ambiguous representations to a stable, category-specific attractor. This dissociation between fixed attractor geometry but feedback-driven trajectory rerouting suggests a general principle for long-range feedback, in which higher cortical regions act as low-dimensional controllers that inject task-relevant priors to guide state-space search of lower cortical regions without requiring relearning of sensory backbone.

**Figure 4.**
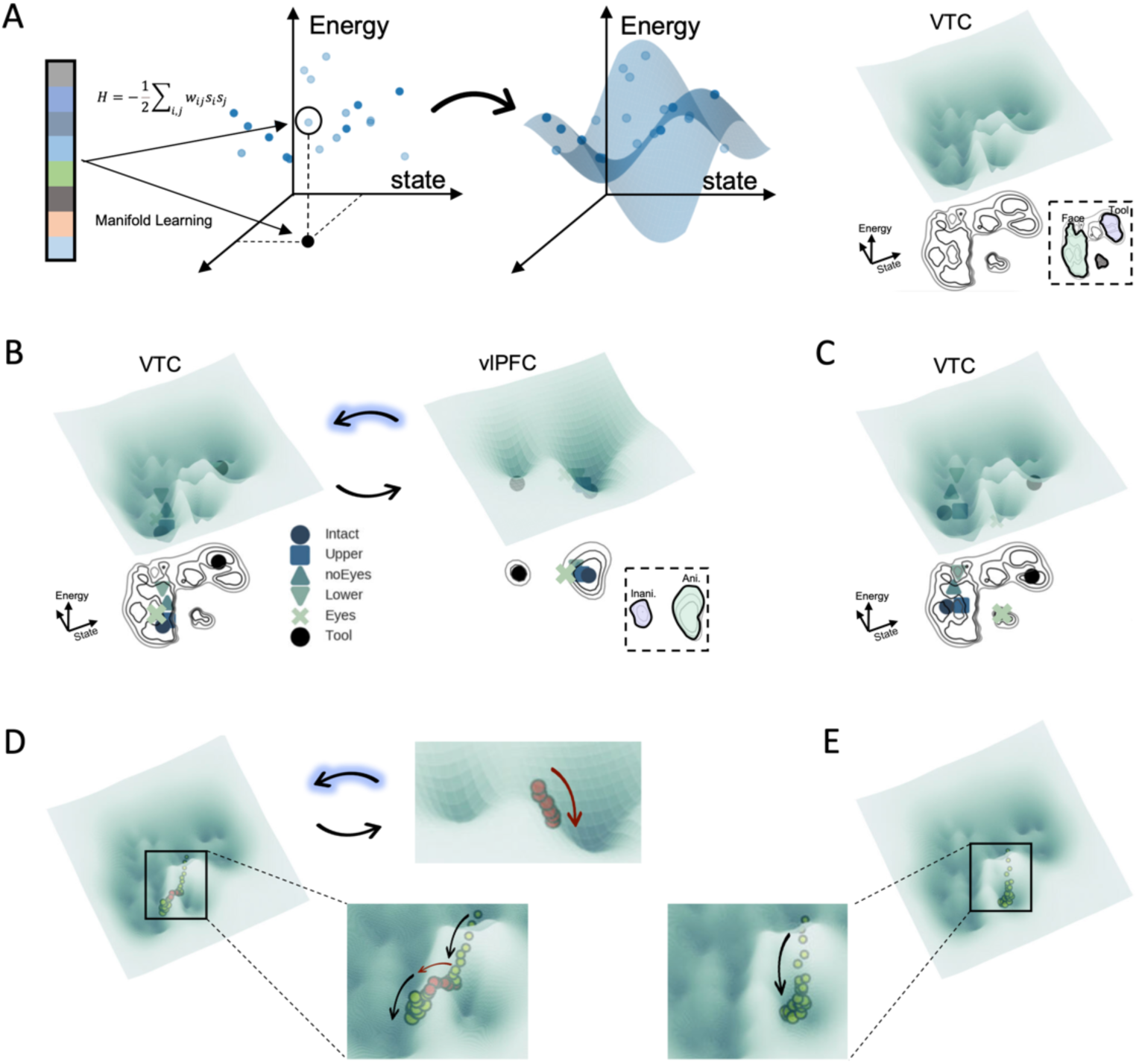
Energy-representation manifold analyses in VTC and vlPFC modules. (A) Left and middle: Schematic illustration of computing energy-representation manifolds. X-Y plane depicts a two-dimensional embedding of high-dimensional VTC neural states, whereas Z-axis represents state energy. Right: the resulting manifold of the VTC module. Light green and purple basins represent faces and tools in the X-Y plane of VTC module, respectively. Note that the basin in gray color indicates a pseudo-state basin corresponding to ambiguous representations. (B) While VTC module exhibits fine-grained category basins, vlPFC manifold is organized along a coarser animacy dimension, separating animate (light green) and inanimate (purple) states. Colored markers indicate neural states under different occlusion conditions. (C) In the absence of vlPFC feedback, eyes stimuli remain trapped in the VTC pseudo-state basin, yielding ambiguous representations. (D) With vlPFC feedback, the state trajectory of severely-occluded faces illustrates an escape from the pseudo-state basin and convergence into the face basin. Yellow dots represent the trajectory of neural states over time within the energy-representation space, while red dots indicate periods involving vlPFC intervention. (E) Without vlPFC feedback, the state trajectory of the same stimulus falls into the pseudo-state basin, whose identity becomes ambiguous.

### EEG reveals delayed responses in processing occluded faces

The energy landscape analysis suggests that engaging vlPFC feedback may incur temporal costs, as additional iterative steps are necessary to resolve ambiguity. Accordingly, the processing of occluded stimuli is expected to require extra time, manifesting as systematic latency shifts in decoding if the brain employs a mechanism analogous to that implemented in the hierarchical vision model. To test this prediction, we used electroencephalography (EEG) capable of high temporal resolution to capture the dynamics of decoding accuracy in a new cohort of participants (N=15) as they viewed occluded faces and tool stimuli from the fMRI experiment. Each stimulus was presented for 500 ms, followed by an interstimulus interval (ISI) of 800–1200 ms (Figure 5A). EEG continuously recorded brain activity, with analyses focused on face-selective channels over the occipitotemporal cortex (Figure 5B; time series for each condition are shown in Supplementary Figure 10), identified via the face-selective N170 component, to examine the temporal processing of occluded faces.

**Figure 5.**
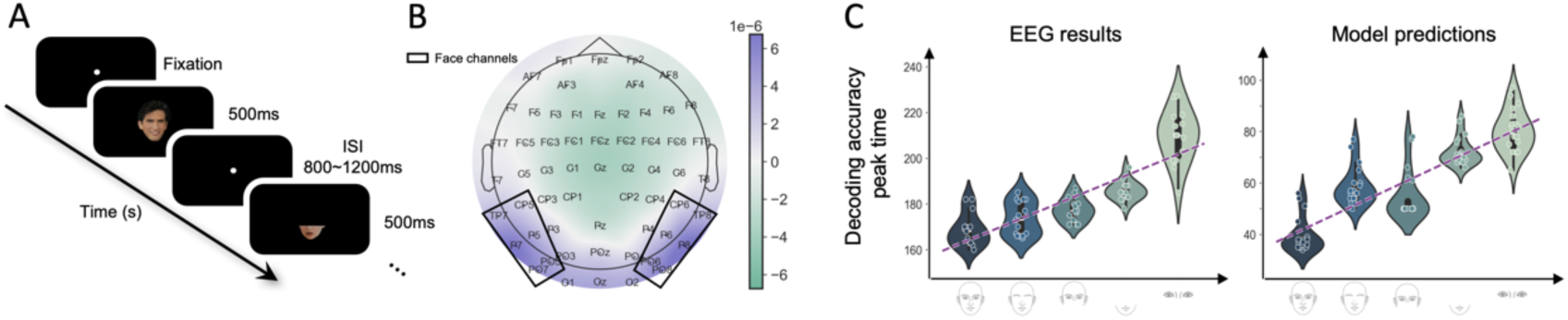
Prolonged latency in decoding occluded faces. (A) Experimental procedure. Stimuli were identical to those used in the fMRI experiment and presented in random order for 500 ms, with ISI ranging from 800 to 1200 ms. (B) EEG channel layout, with black boxes indicating face-selective channels used for analysis. The topographic map shows the average voltage difference between faces and tools across all channels between 165 and 175 ms after stimulus onset, with larger differences indicating greater face selectivity. (C) Peak decoding time for EEG (left) and the hierarchical vision model (right) progressively increased with occlusion intensity.

As expected, we observed the predicted delay in neural decoding under occlusion. Using multivariate pattern analysis of EEG ^42^, we tracked the time course of decoding accuracy in face-selective occipitotemporal channels for each occlusion condition (Supplementary Figure 11A). For intact faces, peak decoding occurred at 170 ms post-stimulus, significantly earlier than for occluded conditions (*t*(14) = –9.78; *p* < .001). Critically, as occlusion severity increased, peak decoding times shifted progressively later (noEyes: 173.60±1.83 ms; Upper: 177.47±1.24 ms; Lower: 185.80±1.07 ms; Eyes: 209.41±2.88 ms; one-way ANOVA across conditions, *F*(4,56) = 68.74, *p* < .001; *R^2^* = 0.80, linear contrast; Figure 5C, left). For comparison, we applied the same decoding procedure to the facial cluster in the hierarchical vision model as its network dynamics evolved (Supplementary Figure 11B), revealing a time-lag effect consistent with EEG findings (*F*(4,95) = 92.54, *p* < .001; *R^2^* = 0.79, linear contrast; Figure 5C, right). Therefore, this graded latency pattern, observed in both the EEG empirical experiment and model simulations, supports the hypothesis that engaging the top-down feedback incurs temporal costs associated with iterative refinement. Such delayed latencies in processing occluded stimuli are consistent with a predictive coding regime, in which initial feedforward representations are insufficient, and additional cycles of feedback are required to achieve a stable perceptual interpretation ^26,70,71^.

Taken together, these findings from fMRI, modeling, and EEG demonstrate that long-range feedback from the vlPFC dynamically modulates representations in VTC, extending processing time to resolve ambiguity.

## Discussion

Occluded recognition transforms vision into an underdetermined inference problem where sparse sensory evidence may support multiple internally plausible interpretations. In this study, we employed systematically graded face occlusion to examine unique contributions of long-range frontal feedback beyond feedforward processing and local recurrence. Integrating findings from fMRI, computational modeling, and EEG, we present a unified mechanistic explanation. Specifically, when sensory evidence for faces became insufficient, the vlPFC sustained a low-dimensional semantic belief state (e.g., animacy vs. inanimacy) and transmitted this abstract information back to the VTC, preferentially targeting the large-scale animacy map encompassing face-selective representations. Critically, in our hierarchical vision model, this frontal feedback operated as a low-dimensional generative input that did not reshape the intrinsic attractor geometry of the VTC; instead, it provided targeted directional input, rerouting ongoing neural dynamics away from an ambiguous state toward a stable, face-consistent attractor basin in the representational energy landscape. This control-based mechanism thus enables perceptual completion in situations where bottom-up sensory signals alone would result in ambiguous representations, and further predicts temporal costs empirically verified through EEG. The predictive attractor-control mechanism identified here in the context of occluded face perception may broadly extend to other perceptual scenarios where the visual system must select among multiple coherent hypotheses under incomplete or degraded inputs, such as clutter, noise, or camouflage, thereby bridging analysis-by-synthesis theories of vision ^34,54^ and dynamical-systems perspectives on the functionality of long-range frontal feedback as state-space control.

A growing body of evidence from primate neurophysiology and human neuroimaging has demonstrated that the frontal cortex contributes to robust object recognition when sensory evidence is incomplete or degraded ^24,25,29,34^. Our study advances this literature beyond the general claim that feedback is beneficial, providing instead a mechanistic and content-specific account. As occlusion increased, the vlPFC did not simply enhance overall visual responsiveness; rather, it provided an abstract semantic constraint that was low-dimensional in structure and selectively routed to the appropriate representational substrate in the VTC. Specifically, decreasing facial information was accompanied by a systematic strengthening of vlPFC-VTC functional coupling in a highly structured manner. This coupling preferentially engages the animacy map, rather than diffusely modulating VTC or merely amplifying activity within the FFA in isolation. This coupling pattern is significant because face representations are embedded within the broader superordinate organization of the animacy map, and delivering an animacy-level prior to this topography therefore constitutes an efficient strategy for biasing inference when bottom-up sensory evidence is compatible with multiple interpretations. Converging evidence regarding this representational content emerges from our geometric analyses. Neural activity in the vlPFC was compressed into fewer dimensions and supported decoding of abstract animacy distinctions more strongly than fine-grained category identity, consistent with the interpretation that the vlPFC conveys a coarse but stable hypothesis, such as animate versus inanimate objects, rather than reconstructing missing visual details at the pixel level. Therefore, our study provides a novel geometry-aware view of predictive coding theories ^34,36,72^. When occlusion pushes ventral visual stream representations into ambiguous attractors of state space, the vlPFC provides a superordinate semantic constraint and directs it to the animacy map organized to exploit that constraint, thereby stabilizing perception without relying on nonspecific attentional gain or feature-level templates.

A central implication of our findings is that frontal feedback functions primarily as a generative constraint rather than as nonspecific amplification of weak sensory evidence. Classical theoretical accounts have long proposed that top-down signals “fill in” missing details when sensory data are incomplete ^10,73^, but such descriptions typically remain general and metaphorical. In contrast, our integrative empirical and modeling approach permits a more precise characterization, specifying the exact form of feedback, elucidating its interaction with VTC representations, and clarifying precisely how frontal feedback resolves perceptual ambiguity. Specifically, frontal feedback did not dynamically reconfigure the VTC circuitry or reshape its attractor geometry. Instead, frontal feedback acts as a low-dimensional control signal, injecting a directed influence along dimensions consistent with animate and specifically face-relevant manifolds. This targeted control reroutes neural trajectories away from ambiguity and into a stable, face-consistent attractor basin. The empirically observed graded latency shifts in the EEG experiment precisely match this proposed mechanism ^6,20,21,42^, as ambiguity resolution relied on iterative correction of neural trajectories rather than instantaneous enhancement of signals. Thus, our combined fMRI, computational modeling, and EEG analyses provide an integrated explanatory framework connecting circuit-level coupling (fMRI), dynamical causal mechanisms (modeling), and the associated temporal costs (EEG), thereby clarifying the inferential role of frontal feedback in perceptual processing.

In summary, our results converge toward a general principle describing the computational role of long-range frontal feedback: high-level semantic knowledge in the vlPFC functions simultaneously as a prior in a Bayesian predictive coding framework and as a targeted control signal in a dynamical-systems context. The key idea of this synthesis lies not merely in acknowledging the general utility of feedback under uncertainty, but in precisely characterizing the nature of its abstract beliefs in high-level cortical regions and its targeted routing toward specific neural attractors in low-level cortical regions. This insight implies that robustness and efficiency in the human visual system may depend less upon continuously improving feedforward invariance, and more upon explicit inference loops mediated by long-range feedback. Therefore, our study points toward a promising direction for developing hierarchical architectures of AI systems, complementing current attention-based and transformer-style integration ^53,74^, and purely reconstruction-based solutions (e.g., image inpainting) ^75^. Rather than primarily scaling up a fast feedforward processing backbone, future architectures may gain robustness and efficiency by incorporating a slow, lightweight, high-level controller that operates in a low-dimensional state space and uses feedback as a targeted control signal over the backbone’s dynamics.

Specifically, such hierarchical architectures would integrate (i) a lower-level module characterized by powerful feedforward and local recurrent connections capable of establishing a rich representational geometry, (ii) a higher-level module tasked with hypothesis maintenance, uncertainty detection, and selective routing, and (iii) recurrent belief-state updates implementing active trajectory correction rather than mere activation gain. AI systems built around this hierarchical controller–backbone loop are more likely to exhibit enhanced sample efficiency and greater resilience to noise, perturbations, and distribution shift. While initially inspired to address visual ambiguity, this hierarchical architecture may effectively generalize to a wide range of functional domains, including cognitive processes (e.g., resolving linguistic ambiguity and context-sensitive comprehension), sensory-motor tasks (e.g., stabilizing motor behaviors in uncertain or dynamic environments), and decision-making contexts (e.g., reasoning, planning, and navigating under conditions of incomplete or ambiguous information).

## Acknowledgments

We thank Yuannan Li for helping collect EEG data. This work was supported by National Natural Science Foundation of China (T2488101), and Beijing Municipal Science & Technology Commission, Administrative Commission of Zhongguancun Science Park (Z221100002722012).

## Author Contributions

YY.Z. conceived the research, collected and analyzed fMRI and EEG data, built the hierarchical vision model, and wrote the article. JR.L. collected and analyzed EEG data, and developed the baseline models. J.L. conceived the research, interpreted the data, and wrote the article.

## Data and code availability

The fMRI and EEG data reported in this paper will be shared by the lead contact upon request. All custom code has been deposited at https://github.com/sdgds/vlPFC-VTC and is publicly available as of the date of publication.

## Materials and Methods

### Generation of Information-Graded Occluded Face Dataset (IGOF)

#### Transfer Learning for Face and Object Representation

To extract high-dimensional feature representations of faces and objects, we employed a transfer learning approach based on the pre-trained AlexNet model. Specifically, we appended a small neural network to the last layer of AlexNet, consisting of two layers: the first layer receives the 1000-unit output from AlexNet and applies a ReLU activation function; the second layer outputs two units for face and object classification, corresponding to face and object recognition, respectively. We trained this transfer learning network using the LFW face dataset ^76^ and the Caltech256 object dataset ^77^. During training, we froze all parameters of AlexNet and only trained the newly added neural network layers using stochastic gradient descent (SGD). This model achieved a classification accuracy of 100% on the test set.

#### Calculation of Face Information in Images

Following transfer learning, we utilized the Grad-CAM technique ^51^ to generate face information heatmaps for each occluded face image. The Grad-CAM method computes the gradients of the input image with respect to the activation of the face unit and combines them with the activation values of the last convolutional layers to produce heatmaps that highlight regions crucial for face feature recognition (exemplars in Supplementary Figure 1). For each occluded face image, we calculated its contribution to the activation of the face unit, thereby obtaining the corresponding heatmap. Subsequently, we averaged the heatmap values to quantify the amount of face information contained in each image. This quantification provided essential quantitative indicators for designing the occluded face stimulus set.

Based on the heatmaps generated by Grad-CAM and the quantified face information, we designed an occluded face stimulus set consisting of 100 images. This set includes five primary occlusion conditions: Intact face, noEyes, Upper, Lower and Eyes, with 20 images per condition. In addition, to compare the brain and model responses elicited by tool images, we introduced an intact tool condition (also consisting of 20 images). All intact images (including intact faces and tool images) were obtained from the Human Connectome Project dataset ^78^. By calculating the heat maps of these images, we manually adjusted the position and area of the occlusion to ensure that the occlusion produced a noticeable gradient change in facial information under different conditions, thereby meeting the requirements of the experimental design.

#### Testing Deep Neural Networks with Occluded Face Stimuli

After creating the occluded face stimulus set, we used it to evaluate the performance of multiple deep neural network models in face recognition tasks. First, we trained the face and object classification units of AlexNet ^52^ (includes other classic DCNNs: VGG19 ^79^, ResNet50 ^80^, ResNet152 ^80^, Inception_v3 ^81^), VIT-B/16 ^5^ and CORnet-S ^3^ models using the transfer learning method described in the previous section. We then tested the classification accuracy of these models under different occlusion conditions with the occluded face stimulus set.

#### fMRI Data Acquisition and Analysis

##### Participants

A total of 30 college students participated in the occluded face fMRI experiment (16 females; mean age = 22.8 years, SD = 2.62 years). All the participants had normal or corrected-to-normal vision, and no history of psychiatric or neurophysiological disorders. The study protocol was approved by the Institutional Review Board of Beijing Normal University, and written informed consent was obtained from all participants prior to the experiment.

##### Stimuli and Experimental Design

This experiment was designed to investigate how the human brain processes partially occluded faces under various masking conditions. We defined six conditions (Intact face, noEyes, Upper, Lower, Eyes and Tools) based on the IGOF dataset. The full IGOF dataset consisted of 20 images per occlusion condition and was used for comprehensive evaluation of model performance. For the fMRI experiments, a representative subset of 10 images per condition was selected from the full stimulus pool to reduce experimental duration and minimize participant fatigue. Subsampled stimuli were balanced across conditions and matched in terms of Grad-CAM–derived face-information levels, ensuring that the critical contrasts between occlusion conditions were preserved. A black dot in the center of the screen was used to mark the fixation point.

In order to maintain participants’ attention throughout the fMRI scanning, a one-back task was implemented. During each run, 70 images were presented in total: the 60 unique stimuli (10 from each condition) plus 10 repeated stimuli randomly interspersed among them. Participants were instructed to press a button if the current image was identical to the immediately preceding one and to refrain from responding otherwise.

The functional scan was acquired using an event-related design with five functional runs. Each trial lasted approximately 5 seconds, encompassing 1 second of stimulus presentation and a variable inter-stimulus interval (3 or 5 seconds) that allowed for jittering. Each run contained 70 trials and lasted about 6 minutes in total, with a 30-second rest interval between consecutive runs. Five runs were administered, leading to an overall scanning time of approximately 33 minutes per participant. A brief 8-second period at the start of each run allowed for MR signal stabilization, and a 12-second period at the end served to mitigate any end-of-run carryover effects. All stimuli were displayed on a projection screen at a resolution of 1024 × 768, with participants positioned approximately 70 cm from the screen to achieve a visual angle of about 6° to 8°. The experimental code was developed in Python and controlled stimulus presentation and response collection via an MRI-compatible button box.

##### fMRI Data Acquisition

All scans were performed on a 3T Siemens MAGNETOM Prisma MRI scanner. Functional images were acquired using a gradient-echo echo-planar imaging (GRE-EPI) sequence with simultaneous multi-slice (SMS) acceleration. The parameters were: repetition time (TR) = 2000 ms, echo time (TE) = 30 ms, flip angle = 90°, field of view (FOV) = 208 × 208 mm, matrix size = 104 × 104, 64 axial slices (2.0 mm thickness), resulting in an isotropic voxel size of 2.0 × 2.0 × 2.0 mm³. For anatomical reference and spatial normalization, a high-resolution T1-weighted structural image was acquired using an MPRAGE sequence. The parameters were: TR/TE/TI = 1500/1.87/756 ms, flip angle = 10°, matrix = 320 × 320, slices per slab = 208, and an isotropic voxel size of 0.8 × 0.8 × 0.8 mm³. Additionally, a gradient echo field map (GRE Field Mapping) was acquired with the following parameters: TR = 566 ms, TE1/TE2 = 4.37/6.83 ms, flip angle = 50°, FOV = 208 × 208 mm, matrix size = 104 × 104, 64 slices, and a voxel size of 2.0 × 2.0 × 2.0 mm³, matching the functional acquisitions for geometric distortion correction.

##### fMRI Data Preprocessing

Functional MRI data were preprocessed using fMRIPrep ^82^. The preprocessing pipeline began with slice timing correction, followed by motion correction through realignment of functional volumes to a reference image. Functional images were then co-registered to each participant’s T1-weighted anatomical scan. Cortical surface reconstruction was performed using FreeSurfer, which involved segmentation of brain tissues and generation of surface meshes. The functional data were projected onto the individual cortical surfaces at the mid-thickness layer and subsequently registered to the fsaverage5 standard surface template to facilitate group-level analyses. Spatial smoothing was applied on the cortical surface using a Gaussian kernel with a 4-mm full-width at half-maximum (FWHM) to enhance the signal-to-noise ratio and account for inter-subject anatomical variability. Additional preprocessing measures included detrending, high-pass filtering (commonly 0.01 Hz), and nuisance regression for motion parameters or physiological signals. Head motion estimates were modeled via the Friston 24-parameter approach ^83^ to mitigate motion artifacts. This unified workflow ensured consistent, high-quality preprocessing of all functional data prior to subsequent statistical analyses.

##### Localization of Individual FFA

To define the individual fusiform face area (FFA), we estimated a first-level general linear model (GLM) using Nilearn (Python). Event onsets and durations were specified from the experimental schedule and modeled as boxcar regressors, convolved with a canonical SPM hemodynamic response function. The model was fit to the concatenated preprocessed fMRI time series using an AR(1) noise model. To reduce nuisance variance, we included as confounds the rigid-body motion parameters, as well as mean signals from cerebrospinal fluid (CSF), white matter (WM), and global signal. The resulting design matrix was used to compute the faces > tools contrast, yielding a vertex-wise *t*-statistic map. We then applied a t-threshold (*t* > 1 in the present implementation) and restricted the surviving vertices within the FFA mask from Zhen et al. ^84^. Finally, the individual right FFA mask was defined.

##### Decoding Occluded Face from FFA

To decode stimulus image category (face vs. tool) from FFA responses, we performed a binary classification analysis using the individually defined FFA as a region of interest (ROI). For each participant, we extracted vertex-wise fMRI responses time-locked to stimulus onset and sampled the response at 2 TRs after stimulus onset (i.e., a canonical peak-latency approximation) to maximize category discriminability.

Because the number of vertices within the individually defined FFA ROI varied across participants, we used a fixed-size feature selection strategy: for each participant, we randomly selected 10 vertices from the individual FFA and computed their response vector for each stimulus. The random vertex selection was performed independently for each participant and repeated across iterations (see below) to ensure that results were not driven by a specific subset of vertices.

For each experimental condition, we first averaged the selected-vertex responses across all trials of the same stimulus within each participant. We then aggregated these stimulus-level response vectors across participants to obtain a group-level FFA response pattern for each stimulus (10-dimensional feature vector; one dimension per selected vertex). This procedure was applied to the intact face and tool conditions to form the training set, and to the occluded-face conditions to form the test set.

We trained a binary face/tool classifier on the group-level FFA response patterns derived from intact faces and tools, and then tested the trained classifier on the group-level response patterns for occluded faces. Importantly, the classifier was identical in type and training objective to the classifier used in the *Transfer Learning for Face and Object Representation* section for the Information-Graded Occluded Face (IGOF) dataset, and the training/testing protocol matched the evaluation framework used in *Testing Deep Neural Networks with Occluded Face Stimuli* section (i.e., a classifier trained on intact face vs. tool and evaluated on occluded-face representations under the same decision rule). To account for stochasticity introduced by random vertex selection, we repeated the entire sampling–feature construction–training–testing pipeline 30 times and reported the mean classification accuracy across repetitions as the final decoding performance.

##### Surface-based Functional Connectivity Analysis

In order to locate the neural top-down origins of the modulated FFA signals, we conducted the surface-based searchlight analysis method. First, we defined the ventrolateral prefrontal cortex (vlPFC) ROI based on the HCP-MMP parcellation. Specifically, the vlPFC ROI comprised parcels 6r and IFJp in the right hemisphere, and FOP4 and FOP1 in the left hemisphere ^55^. Second, a searchlight radius of 5-mm was set on the fsaverage5 standard surface to find the neighboring vertices of each vertex within the set radius. For each participant, activation values of vertices in vlPFC were first extracted for the second TR after all stimulus occurrences in a given condition. These activation values represented the immediate brain response to stimulus processing. Subsequently, the Euclidean distance matrix between vertex activation values within the defined individualized FFA region across stimuli was computed, and the same calculation was repeated for the set of activation values within a 5-mm radius around each vertex in the vlPFC. Based on these distance matrices, correlation coefficients between FFA and vlPFC vertices were calculated as an indicator of the strength of functional connectivity between them. In this way, we were able to assess the patterns of functional connectivity between the FFA and vlPFC vertices across the brain.

For each participant and each occlusion condition, we obtained a vertex-wise functional-connectivity map within the vlPFC ROI (defined by HCP-MMP parcels; see ROI definition above) by computing the correlation coefficient (r) between the FFA distance matrix and the distance matrix of the 5-mm searchlight centered at each vlPFC vertex. To summarize these connectivity maps, we converted r values to two-tailed significance values and applied a Bonferroni correction for multiple comparisons across vertices within the vlPFC ROI. Vertices surviving the corrected threshold were labeled as significantly connected to FFA for that condition. We then computed two summary measures per condition: (i) the proportion of significant vlPFC vertices and (ii) the mean connectivity strength, defined as the mean r value across significant vertices. These two measures correspond to the left and right panels in Figure 2B. At the group level, we averaged each measure across participants for each occlusion condition and performed a linear regression with occlusion severity as the predictor (5 levels) to test whether vlPFC–FFA coupling increased monotonically with occlusion, as reported in Figure 2B.

To evaluate whether the occlusion-dependent vlPFC–FFA connectivity effects were specific to the initially selected vlPFC parcels, we conducted a control analysis by repeating the same searchlight connectivity procedure within an a priori LPFC search space defined using HCP-MMP parcels associated with lateral prefrontal cortex (Supplementary Figure 6 top). For each participant and each occlusion condition, we computed vertex-wise connectivity (r) between the FFA distance matrix and the searchlight distance matrix centered at each LPFC vertex (5-mm radius; identical to the primary analysis). Within each occlusion condition, we identified LPFC vertices showing significant connectivity with FFA after Bonferroni correction across vertices. We then aggregated results at the parcel level: for each LPFC parcel, we calculated the mean r across significant vertices within that parcel for each condition. For each parcel, we performed a linear regression across the five occlusion conditions on the group-averaged parcel connectivity and extracted the *p* value associated with the slope term. Parcel-wise *p* values were Bonferroni-corrected across LPFC parcels, and parcels surviving the corrected threshold (*p* < .05) are shown in black in Supplementary Figure 6 bottom, with the remaining parcels colored by their *p* values.

To investigate the neural mechanisms that regulate FFA signal processing from the top down, we performed a surface-based searchlight analysis to compute functional connectivity between the ventrolateral prefrontal cortex (vlPFC) and the ventral temporal cortex (VTC). The analysis was conducted on the fsaverage5 surface model with a 5-mm searchlight radius. We defined a human VTC template using the HCP-MMP parcellation by combining eight parcels (V8, FFC, PIT, PHA3, TE2p, PH, VMV3 and VVC) into a single coherent region ^85,86^. Within this VTC template, we further partitioned vertices into an animacy map and an inanimacy map using the mid-fusiform sulcus (MFS) as an anatomical boundary. The MFS is a shallow longitudinal sulcus that reliably divides the fusiform gyrus into lateral and medial partitions and has been shown to align with large-scale functional divisions in human VTC ^85,87^. Consistent with prior work linking this landmark to category-related functional topographies, we treated vertices lateral to the MFS as belonging to the animacy-related sector and vertices medial to the MFS as belonging to the inanimacy-related sector (i.e., animacy and inanimacy maps, respectively).

The functional-connectivity computation followed the same procedure described above, except that we used vlPFC as the seed and VTC vertices as targets. Specifically, for each occlusion condition, we computed the distance matrix of stimulus-evoked response patterns within the vlPFC seed and correlated it with the distance matrix computed within the 5-mm searchlight centered on each VTC vertex, yielding a vertex-wise connectivity map. We then quantified connectivity selectively within the animacy and inanimacy maps to assess whether vlPFC coupling preferentially targeted VTC maps implicated in abstract representations of animacy versus inanimacy.

##### Effective Dimensions and Radii

Chung and colleagues ^56^ developed a theory of linear divisibility of manifolds, which defines the effective dimension and radius of a single manifold using anchors at the edges of the manifold. Specifically, the effective dimension is defined as the spread of these anchor points along the different axes of the manifold, while the radius is the total variance of the anchor points normalized by the mean distance from the center of the manifold. To apply Chung et al.’s method to our data, we first randomly selected 100 vertices in each of the VTC and vlPFC. We then applied Chung et al.’s method to calculate the effective dimension and radius of each manifold. This procedure was repeated 20 times to obtain reliable values for the VTC and vlPFC.

##### Neural Decoding Abstract Information

To test the hypothesis that vlPFC preferentially encodes abstract information, we conducted multivariate decoding analyses in vlPFC and VTC using Haxby object-vision dataset ^64^. The central comparison was between decoding at an abstract level (animacy vs. inanimacy) and at a fine-grained level (specific object category). Abstract labels were defined as animate (faces and cats) versus inanimate (houses, bottles, scissors, shoes, and chairs), and fine-grained decoding treated the same seven categories as distinct class labels. Decoding performance was compared between vlPFC and VTC to assess whether abstract category information is more strongly represented in vlPFC than in VTC.

For the Haxby dataset, we extracted stimulus-evoked multivoxel response patterns from each ROI across the full time series after excluding rest periods. Thus, each sample corresponded to an fMRI volume acquired during object viewing, represented as a vector of voxel responses within the ROI. ROI masks were defined using the HCP-MMP atlas labels in volumetric space to obtain vlPFC and VTC patterns in each hemisphere. The vlPFC and VTC templates utilized the same brain regions as “The Surface-based Functional Connectivity Analysis” section. These ROI pattern vectors were paired with the corresponding category labels (face, cat, house, bottle, scissors, shoes, chair) to form the training and testing data for decoding.

Decoding was performed using a linear support vector machine (SVM). To minimize sensitivity to any particular subset of voxels and to obtain a stable estimate of information content, we employed a repeated feature-subsampling procedure: on each repetition, we randomly selected a fixed number of voxels from the ROI and evaluated classification accuracy, yielding a distribution of accuracies rather than a single point estimate. Classifiers were trained on a subset of samples and tested on held-out samples that were not used for training. Abstract decoding used binary labels (animacy vs. inanimacy), whereas fine-grained decoding used a 7-way classifier.

To enable direct comparison between abstract and fine-grained decoding despite their different chance levels, decoding performance was reported as chance-corrected accuracy (observed accuracy minus 1/2 for abstract decoding and minus 1/7 for fine-grained decoding). Finally, we statistically compared abstract versus fine-grained decoding within each ROI and tested whether the abstract–fine advantage was larger in vlPFC than in VTC.

#### Building the vlPFC-VTC model

The vlPFC-VTC model contained two coupled subsystems. (i) A “ventral visual module” that transformed images into a topographic cortical sheet representation. This part followed our previous work, where a pre-trained AlexNet provided high-dimensional image features and a self-organizing map (SOM) learned a two-dimensional VTC-like organization via competitive learning and neighborhood-based weight updates. In our subsequent work, this VTC sheet was further endowed with recurrent (Hopfield-like) dynamics and structured interconnectivity governed by an exponential distance rule (EDR), yielding stable attractor-like category patterns. This model was shown to resemble VTC in both function and functional organization ^57,86^. (ii) A vlPFC recurrent sheet that received feedforward signals from VTC and sent low-dimensional, abstract feedback to VTC. Its construction was constrained by our fMRI results but its learning rules were deliberately chosen to remain mathematically consistent with the “ventral visual module”.

##### Model Architecture and Dynamics

The model consisted of the following computational components:

(i) DCNN encoder. A pre-trained AlexNet encoded each image into a feature representation, which served as the bottom-up drive for the VTC module. This network provided high-dimensional object-space features for subsequent cortical-sheet learning.
(ii) VTC module. The VTC was implemented as a two-dimensional lattice (200×200). Its internal dynamics followed a Hopfield-like update rule in which neurons took binary states (+1/−1) and were asynchronously updated by integrating recurrent input and external fields (bottom-up from DCNN, and top-down from vlPFC). This VTC module integrated biological constraints with functional learning principles through a two-stage architecture. In the first stage, visual stimuli were encoded into high-dimensional feature vectors using the DCNN encoder. These features were projected onto a two-dimensional grid via a self-organizing map (SOM), which formed category-specific clusters, mirroring the functional specialization observed in the human VTC ^86^. In the second stage, spatially constrained lateral connections were incorporated into the SOM using an exponential distance rule (EDR), where connection probabilities decayed exponentially with Euclidean distance between neurons. Synaptic plasticity was governed by Hebbian and anti-Hebbian rules: excitatory connections within clusters were strengthened to enhance category-specific representations, while inhibitory cross-cluster interactions suppressed conflicting activations ^57^. This architecture was shown to systematically replicate various neural representations and functional organizations of VTC.
(iii) vlPFC module. The vlPFC was implemented as a smaller two-dimensional lattice (20×20) to represent lower-dimensional abstract structure. Like VTC, vlPFC neurons were binary and evolved according to recurrent Hopfield-like dynamics driven by feedforward input from VTC and its own recurrent connectivity.

Neurons in the VTC and vlPFC modules shared similar architectures and dynamics, updating their states via nonlinear activation of external and internal inputs. The dynamics of VTC neurons were formalized as:

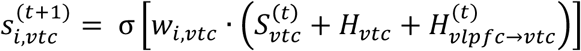

where 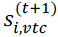 denoted the state of the neuron *i* in the VTC at the next time. The nonlinear activation function was defined as σ(x) = +1 if x ≥ 0, and σ(x) = −1 otherwise. Here, *w_i,vtc_* denoted the vector of recurrent weights projecting to neuron *i* in the VTC, 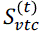 denoted the population state at time *t*, *H_vtc_* denoted the bottom-up input from the DCNN, and 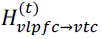 denoted the top-down neural input from vlPFC at time *t*. The vlPFC output signals were passed back to the VTC through the feedback connections to regulate the activity patterns of VTC neurons.

Specifically, this process was expressed by the following equation:

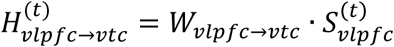

where 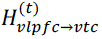 denoted the top-down signal at time *t*, *w_vlpfc→vtc_* denoted the connection from vlPFC to VTC, and was the transpose 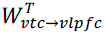 of the feedforward connection from VTC to vlPFC.

The dynamics of vlPFC neurons were defined analogously. In vlPFC, neuronal states were updated based on recurrent input and feedforward input from VTC. The process was represented by the following mathematical expression:

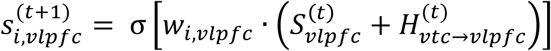

where 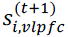 denoted the neural state of the vlPFC at the next time, *w_i,vlpfc_* denoted the weight vector of recurrent connections of all neurons within the vlPFC to the *i*-th neuron, 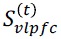 denoted the activation pattern of the neurons in the population of the vlPFC at time *t*, 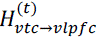 denoted the bottom-up neural input provided by VTC to vlPFC at time *t*:

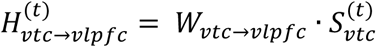

where *w_vtc→vlpfc_* denoted the feedforward connection from the VTC to the vlPFC, and 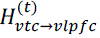 denoted the feedforward signal received by vlPFC from VTC at *t*.

##### Network Learning

Because “learning” in this model referred to multiple weight matrices trained under different rules, we specified each set separately below. These rules were chosen to preserve continuity with our earlier frameworks.

(i) VTC recurrent weights. All VTC-internal parameters followed the same settings as our previous VTC recurrent model ^57^. VTC recurrent connectivity was constructed by Hebbian-style pattern storage and was structured by a distance-dependent rule so that lateral connections obeyed biologically motivated wiring constraints.
(ii) VTC-to-vlPFC feedforward weights and the feedback weights. To maintain a unified learning principle between “DCNN→VTC” and “VTC→vlPFC”, we trained these weights using the same competitive learning mechanism used in SOM formation: at each iteration, the current VTC population state was treated as an input vector; the closest (winner) vlPFC unit was selected; and the winner plus its neighbors updated their weight vectors according to a Gaussian neighborhood function. This belonged to the same algorithmic family as the SOM learning rule described in our “ventral visual module”, where neighborhood-based updates enforced smooth topography on a cortical sheet. We used the validation dataset of ImageNet 2012 ^88^ with all 50,000 images for a total of 100,000 iterations to train *W_vtc→vlpfc_*. The top-down feedback weights *W_vlpfc→vtc_* were not learned independently. Instead, we defined them as proportional to the transpose 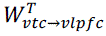 of the feedforward connection from VTC to vlPFC. This choice mathematically linked the bidirectional pathways, effectively reduced the number of free parameters and ensured that the observed top-down effects emerged from the system’s inherent recurrent dynamics rather than from additional parameter fitting. Biologically, this bidirectional coupling was a reasonable abstraction because reciprocal feedforward–feedback projections are pervasive between cortical areas in the primate visual hierarchy, consistent with the widespread existence of paired inter-areal forward and feedback pathways ^1,2^. Motivated by anatomical evidence that top–down inputs can constitute a substantial fraction of synaptic drive relative to feedforward afferents (often differing by several-fold, around 3 to 6-fold, in classic visual relay circuits) ^89,90^, and by temporal evidence that prefrontal top–down influences primarily emerge after the initial feedforward sweep (approximately 130 ms) and shape late-phase ventral-stream representations ^34^, we implemented a delayed, high-gain vlPFC-to-VTC feedback term and set its gain to four times that of the VTC-to-vlPFC feedforward 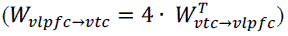 drive as a parsimonious setting within this biologically plausible several-fold regime.
(iii) vlPFC recurrent weights. Our fMRI results showed that vlPFC encoded more abstract representations than the VTC. Accordingly, a key requirement of the vlPFC–VTC model was that vlPFC learn abstract conceptual representations internally. We did this by learning the appropriate weights *W_vlpfc_* in vlPFC.

We then followed the method of Zhang et al. ^57^ to train *W_vlpfc_*. We trained vlPFC recurrent connectivity using the same Hopfield-style learning pipeline as our recurrent VTC model: weights were first built by a Hebbian outer-product rule from target memory patterns, and then constrained by a spatial kernel to impose structured connectivity. To enable *W_vlpfc_* to encode abstract concepts, we trained it to learn both animate and inanimate states as attractors. To do this, we first needed the animate and inanimate states in the vlPFC module, which was done by feeding the model with animate and inanimate pictures after the previous training step to determine the corresponding activation patterns in vlPFC, which were used as memory patterns to learn *W_vlpfc_*. We used 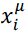 to denote the memory pattern of the *i*-th neuron at the *µ*-th pattern.

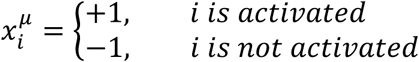

*W_vlpfc_* was established based on Hebbian and anti-Hebbian learning rules, which allowed the network to strengthen or weaken connections depending on the correlated activity of the neurons. Each element in *W_vlpfc_* was updated by:

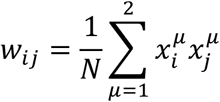

where *N* denoted the total number of neurons, *W_ij_* denoted the connection weight between neurons *i* and *j*, and is the element in row *i* and column *j* of *W_vlpfc_* Self-connections were set to zero. This pipeline ensured that the vlPFC-VTC model was both mechanistically interpretable (each weight set has a clear origin and purpose) and methodologically consistent with the two preceding models it built upon.

##### Model Runs

The model was simulated in two configurations: (i) a baseline configuration without vlPFC, in which the VTC module was driven only by bottom-up input and its intrinsic recurrent dynamics; and (ii) a full configuration that additionally included vlPFC and top-down feedback to VTC.

In the baseline configuration, each image was first processed by the DCNN to obtain a bottom-up input vector to VTC. This input was normalized and stochastically sampled to initialize the binary neuronal states of VTC. Simulations proceeded in two sequential phases. Phase I consisted of 50,000 update iterations during which the bottom-up input remained clamped while VTC states were updated under recurrent dynamics. Phase II consisted of 80,000 further iterations in which the bottom-up input was removed (set to zero), and VTC activity evolved solely under recurrent dynamics.

In the full configuration, Phase I and Phase II were identical to the baseline configuration, except that an additional feedback phase was inserted immediately after Phase I. During this feedback phase (30,000 iterations), vlPFC received the current VTC state via feedforward projections, updated its own state under recurrent dynamics, and then delivered top-down feedback to VTC, which was integrated into the VTC update rule.

##### Decoding Faces and Tools

To quantify category information in VTC activity, we trained a binary classifier (consistent with *Transfer Learning for Face and Object Representation* section) to discriminate faces vs. tools using VTC responses to intact face images and tool images. For each training run, we randomly selected 1,000 neurons from the VTC lattice and used their responses as features. The trained classifier was then evaluated on VTC responses evoked by occluded face stimuli, yielding a classification accuracy for occluded-face representations under the same decision boundary. To account for variability due to random neuron subsampling, the entire training–testing procedure was repeated 30 times with independent random selections of 1,000 VTC neurons, and results were reported as the mean classification accuracy across repetitions (with variability estimated across repeats).

##### Calculation of the Representational Geometry

To quantify the geometric structure of population representations in VTC and vlPFC, we computed the effective dimension and radius of each representational manifold following the framework of Chung and colleagues ^56^. To apply this method to our model representations, we first randomly sampled 100 neurons from each module (VTC and vlPFC) and formed population response vectors in this reduced subspace. We then applied Chung et al.’s procedure to estimate the effective dimension and radius for each manifold in each module. To obtain stable estimates that were not driven by a particular neuron subset, we repeated the random-neuron sampling and the full manifold-geometry computation 20 times, and reported results averaged across repetitions (with variability quantified across repeats).

##### Energy-representation Manifold

The method for calculating three-dimensional energy-characterized manifolds comes from the study by Zhang et al. ^57^. First, we used the UMAP method ^91^ to reduce the dimensionality of the model’s representation of all IGOF stimuli to a two-dimensional space, obtaining a low-dimensional representation that preserves the structure between the high-dimensional representations. Second, since the model is a Hopfield-like neural network, the states of the model evolve towards states with smaller energy, reflecting the model’s tendency to minimize its energy ^92^. To depict this dynamic characteristic, we calculated the energy for each state of the network from the initial state to the stable state using the following equation:

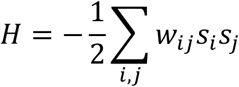

where *H* denoted the energy as a function of the network’s state, *s_i_* and *s_j_* are the states of neuron *i* and *j*, respectively, and *W_ij_* is the connection weights from neuron *j* to *i* ^92^. We incorporate these energy values as a third axis, along with the two-dimensional representation space obtained from the parameterized UMAP, and use interpolation to obtain a smooth surface, thus creating what we call the energy representation space ^57^.

##### Conditional-GAN and Quantitative Analysis of its Generated images

To visualize the information contained in the model’s VTC states, we trained a class-conditional generative adversarial network (cGAN) ^67^ to reconstruct stimulus images from high-level CNN feature vectors. The generator received a feature representation together with a category label and produced an RGB image consistent with these inputs, following the standard conditional GAN framework. The model was trained on a combined dataset of Caltech-256 ^77^ and LFW ^76^, spanning both faces and non-face objects. The generator–discriminator pair was optimized using a least-squares adversarial objective (LSGAN) ^93^ together with a perceptual loss computed on multi-layer VGG features ^94^, encouraging reconstructions that are both visually realistic and perceptually faithful. To decode images from the vlPFC-VTC model, we converted VTC module states into the same 1000-dimensional AlexNet feature format used to train the cGAN. Concretely, we inversely transformed the state of the VTC module along the network architecture to the 1000-dimensional representation, which was then directly provided as input to the trained cGAN to generate a reconstructed image. Using this pipeline, we reconstructed images from the VTC representations for all stimulus conditions from models with and without the vlPFC module.

To quantify how much face-specific content was present in reconstructed images, we computed a face-information score for each generated image using the same procedure described in *Calculation of Face Information in Images* section. We then used this score to quantify restored face information under occlusion. Specifically, the degree of restoration was calculated as the difference between the face information scores in the reconstructed image in the final and initial states of the VTC model. This restored face information was compared between model variants with vs. without vlPFC, providing a quantitative measure of how vlPFC feedback enhances the recovery of face-related content in the generative reconstructions. Furthermore, to quantitatively assess the fidelity of images generated by models with and without the vlPFC module’s steady-state representations, we computed two complementary similarity metrics between the reconstructed images and the ground-truth stimuli. The Structural Similarity Index (SSIM) ^95^ was employed to assess perceived similarity in brightness, contrast, and structural information at a global level. Furthermore, we utilized the Learned Perceptual Image Patch Similarity (LPIPS) metric ^96^, which leverages features from pre-trained deep neural networks to measure similarity in high-level feature spaces, thereby providing evaluations aligned with human perceptual judgments. Higher SSIM and 1-LPIPS (as employed here) scores indicate superior reconstruction quality. The combined use of these metrics delivers a comprehensive evaluation, ensuring generated images remain faithful to the original stimulus in both low-level structural properties and high-level perceptual features.

#### ERP Experiment and Data Analysis

##### Participants

Fifteen healthy volunteers (8 males, 7 females, aged 22–30 years) participated in the study. All participants had normal or corrected-to-normal vision and no history of prosopagnosia, with intact face recognition abilities. EEG data were recorded using a 64-channel wireless BrainCo system. Participants performed tasks in a soundproof, dimly lit EEG chamber to minimize external noise and light interference.

##### Stimuli and Procedure

The experiment utilized a visual paradigm with a one-back task to maintain participants’ attention. Stimuli comprised six conditions: Intact face, noEyes, Upper, Lower, Eyes, and tools. All face and occluded face stimuli were consistent with those used in the fMRI experiment. Each condition included 10 unique stimuli, totaling 60 stimuli. For each experimental run, 10 stimuli were randomly selected for the one-back task, resulting in 70 trials per run (including repetitions). In the one-back task, participants pressed a key if the current stimulus matched the previous one and withheld response otherwise. Stimuli were presented in random order for 500 ms, with inter-stimulus intervals uniformly distributed between 800 and 1200 ms. Each participant completed six runs. The experimental protocol was implemented using the PsychoPy package in Python.

##### Data Preprocessing

EEG data preprocessing involved the following steps: channel localization, selection of all 64 channels, rejection of bad channels (none were excluded), application of a 50 Hz notch filter to remove powerline interference, and a 1–40 Hz bandpass filter to focus on relevant frequency bands. Events were segmented from 100 ms pre-stimulus to 250 ms post-stimulus, followed by baseline correction using the pre-stimulus interval. All preprocessing was performed using the EEGLAB toolbox in MATLAB.

##### Selection of Face-Selective Channels

To identify face-selective channels, preprocessed EEG data were averaged across participants to obtain mean time series for each channel. Channels were deemed face-selective based on two criteria: (i) a pronounced negative deflection (N170) at approximately 170 ms post-stimulus in the intact-face condition; and (ii) a negative difference in mean amplitude between the intact-face and tool conditions within the 165–175 ms window, indicating the strongest N170 for faces. Selected face-selective channels included TP7, P7, PO7, CP5, P5, PO5 (left hemisphere) and TP8, P8, PO8, CP6, P6, PO6 (right hemisphere). These analyses were conducted using the MNE package in Python.

##### Multivariate Pattern Analysis (MVPA) Decoding for EEG and Computational Model

To characterize the temporal dynamics of category information, we performed time-resolved pairwise decoding between each face condition and the tool condition using a linear support vector machine (SVM). For each participant, epoched single-trial data were organized as a tensor of trials × channels × time, restricted to the face-channel set described above. Decoding was performed independently at each time point from 100 to 250 ms post-stimulus. At each time point, we implemented a repeated leave-one-trial-per-class-out procedure. In each iteration, trials were randomly permuted within each class; all but one trial from each class were used for training, and the remaining one trial per class was held out for testing. An SVM classifier was trained on the concatenated training set and evaluated on the held-out test pair. This procedure was repeated 100 times per time point, and decoding accuracy was averaged across repetitions to yield a robust estimate of time-resolved decoding performance for each participant.

To enable a comparison with EEG, we applied the time-resolved MVPA decoding procedure to the model’s VTC dynamics. For each stimulus condition, we extracted trial-wise VTC activity across recurrent processing steps (20 trials per condition). Analyses were restricted to units within the model’s face-selective region. In each repetition, we randomly sampled 100 units from this region and constructed, at each time step, a feature matrix of trials × units for each condition. Pairwise decoding (each face condition vs. tools) was performed independently at each time step using SVM. Following the EEG analysis, we used a repeated leave-one-trial-per-class-out scheme. This procedure was repeated 100 times with random reassignment of held-out trials, and accuracies were averaged to yield the decoding accuracy for that time step. To reduce dependence on the particular subset of sampled units, the entire decoding pipeline was repeated 20 times with independently sampled unit subsets.

## Supplementary Information

**Supplementary Figure 1.**
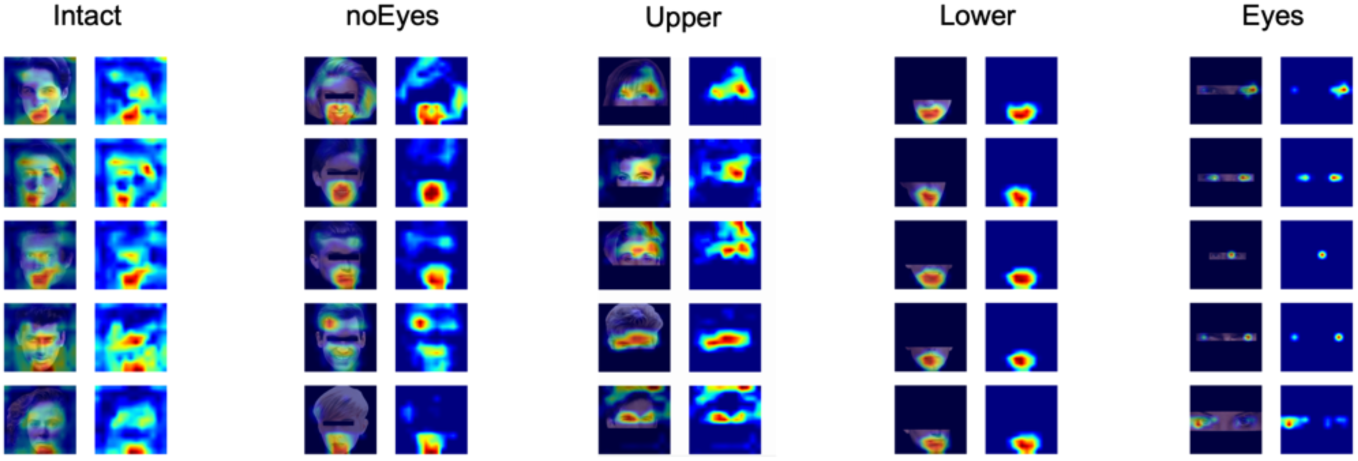
Grad-CAM heatmaps of five different samples under different occlusion conditions. In each sample, the right column shows the heatmap of the face unit, and the left column shows the overlay of the heatmap and the original stimulus.

**Supplementary Figure 2.**
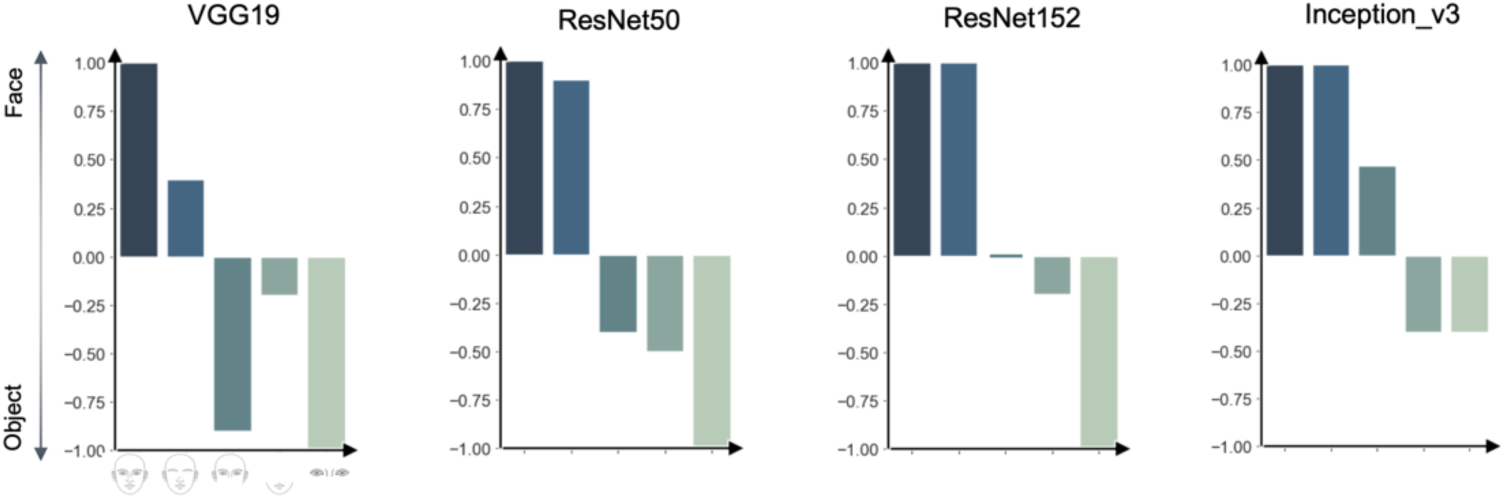
The performance of four other classic visual recognition feedforward convolutional neural networks on the IGOF dataset. The responses and accuracy rates of these feedforward networks exhibit varying degrees of vulnerability to occluded faces.

**Supplementary Figure 3.**
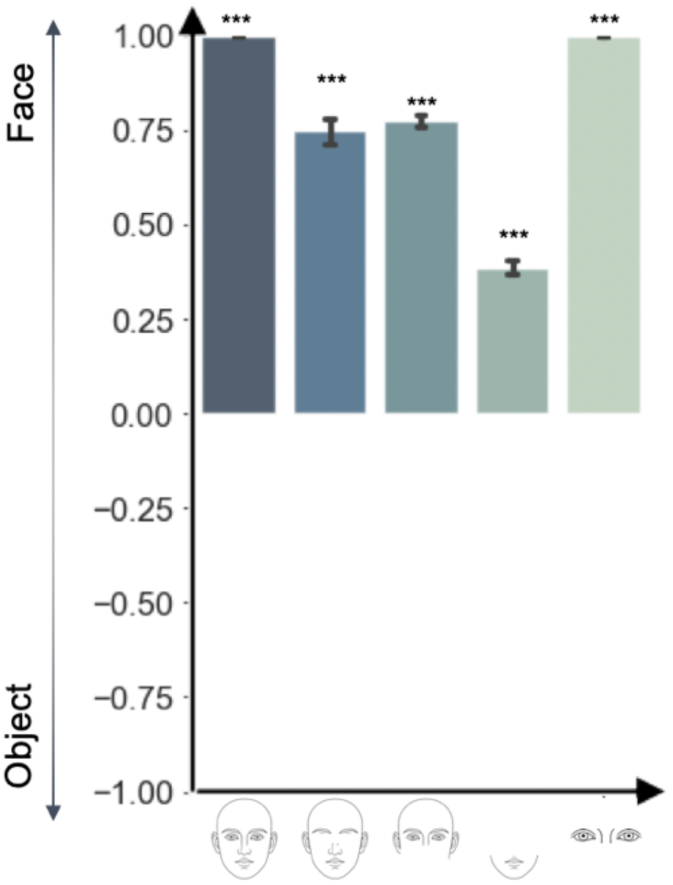
The resilience of right FFA decoding performance across all occlusion levels. Decoding analysis of the fMRI data revealed that the right FFA reliably distinguished faces from non-face objects across conditions (*p*s < .001, Bonferroni corrected).

**Supplementary Figure 4.**
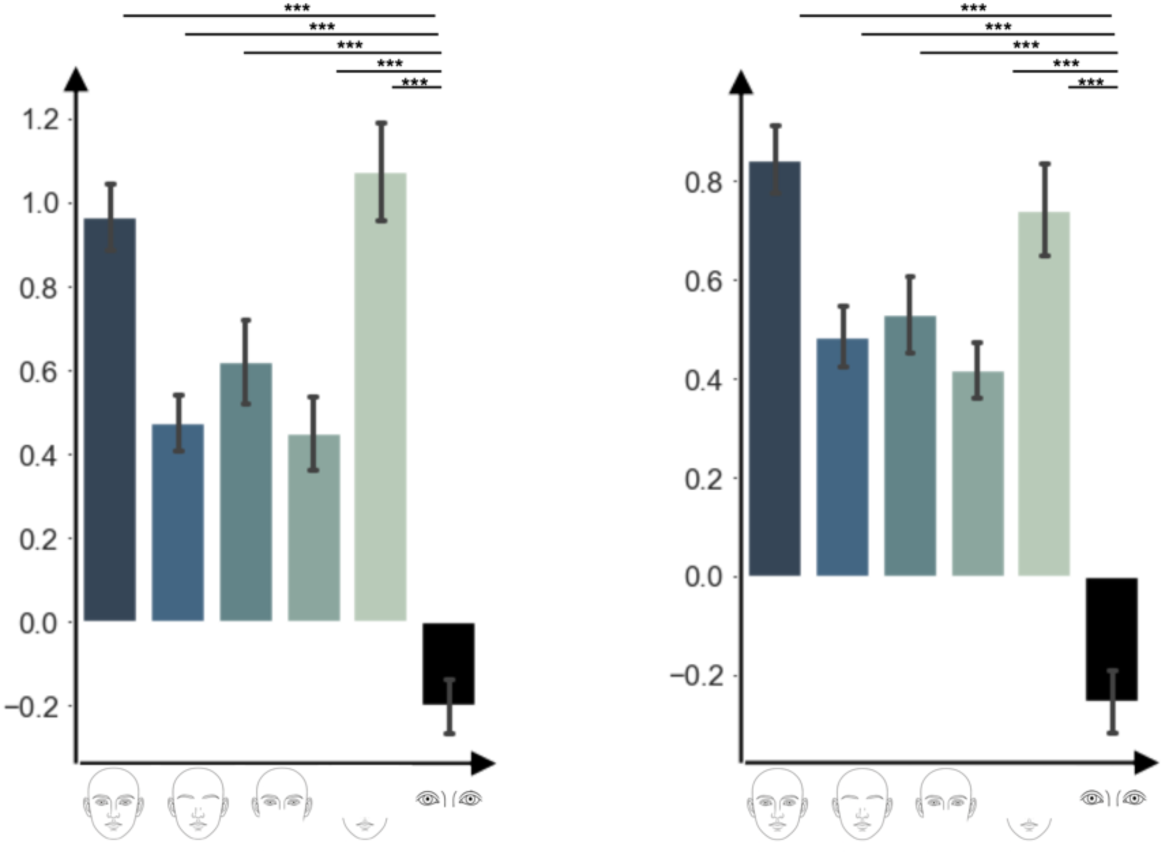
The resilience of FFA GLM analysis across all occlusion levels. GLM analysis of the fMRI data revealed that the beta values in FFA reliably distinguished faces from non-face objects across conditions (*p*s < .001, Bonferroni corrected) in both left (left panel) and right (right panel) hemispheres.

**Supplementary Figure 5.**
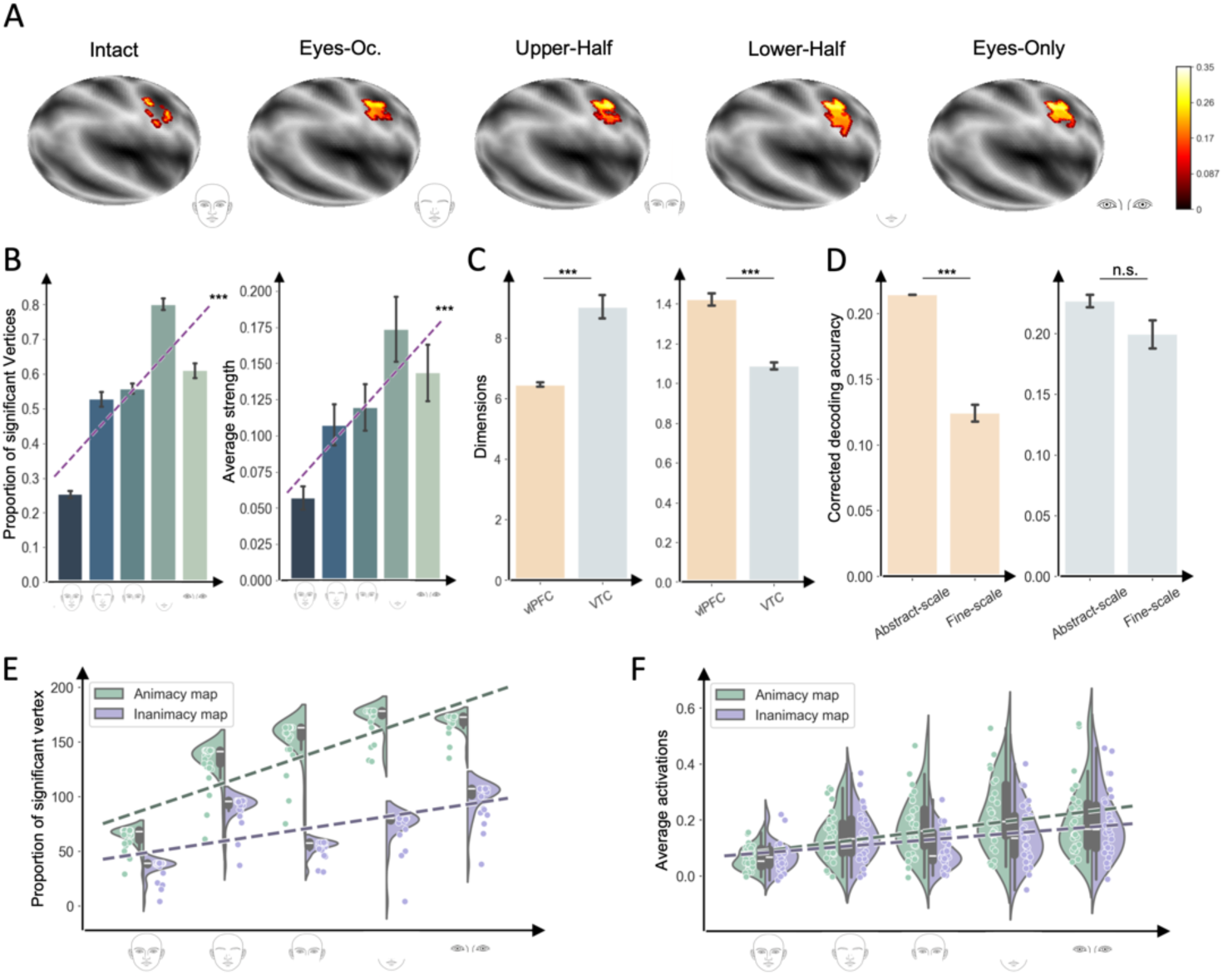
The vlPFC in the right hemisphere provides feedback regulation of the animacy map in the VTC. (A) Cortical vertices in the vlPFC show significant functional connectivity with the FFA under different degrees of occlusion. Dark and light gray indicate sulci and gyri, respectively; color bars represent connectivity strength. (B) Changes in the proportion of cortical vertices (left) and connectivity strength (right) in the vlPFC that are functionally connected to the animacy map under different degrees of occlusion. Linear regression analysis showed significant correlations (proportion of vertices: *R²* = 0.78, *p* < .001; connectivity strength: *R²* = 0.16, *p* < .001), indicating that the vlPFC-VTC coupling increases with increasing occlusion. (C) Effective dimension (left) and representation radius (right) of the geometric manifold in the vlPFC and VTC. vlPFC representations had significantly lower effective dimensionality than VTC representations (vlPFC: 6.46 dimensions; VTC: 9.03 dimensions; *t*(29) = –6.63, *p* < .001), suggesting that the vlPFC compresses sensory information into fewer, more abstract dimensions. Additionally, the representational radius, which reflects the overall span of representations, was significantly larger in the vlPFC than in the VTC (vlPFC: 1.42; VTC: 1.09, arbitrary units; *t*(29) = 9.65, *p* < .001), suggesting that vlPFC representations are more generalized, encoding a broader range of stimuli than VTC representations. (D) Multivariate decoding analysis showed that vlPFC encoded abstract information better than encoding individual categories (left panel), while VTC did not show a preference (right panel). Error bars represent the standard error (SEM) of the mean. A significant region and scale interaction for decoding accuracy (two-way ANOVA *F*(1, 76) = 25.62, *p* < .001) indicates that differences in decoding accuracy between abstract and individual identity scales vary across brain regions. Furthermore, within the vlPFC, we found that vlPFC activity enabled significantly better decoding of abstract categories than individual identity categories (*p* < .001, Tukey HSD). In contrast, VTC decoding accuracy showed no preference for abstract categories. These findings suggest that the vlPFC preferentially supports abstract representations, while the VTC performs equally well across all scales. (E) With the reduction of facial information, the proportion of vertices of functional connections in the animacy map significantly increased (*F*(1,28) = 11.55, *p* < .05; *R²* = 0.79), while the change in the inanimacy map was not significant (*F*(1,28) = 2.50, *p* = .21; *R²* = 0.46). (F) With increasing occlusion, a significant increase in connectivity with the ventrolateral prefrontal cortex (vlPFC) was observed only in the animacy map (*F*(1,28) = 14.40, *p* < .05; *R²* = 0.83), while this phenomenon was not observed in the inanimacy map (*F*(1,28) = 5.34, *p* = .10; *R²* = 0.64). ***: p < 0.001

**Supplementary Figure 6.**
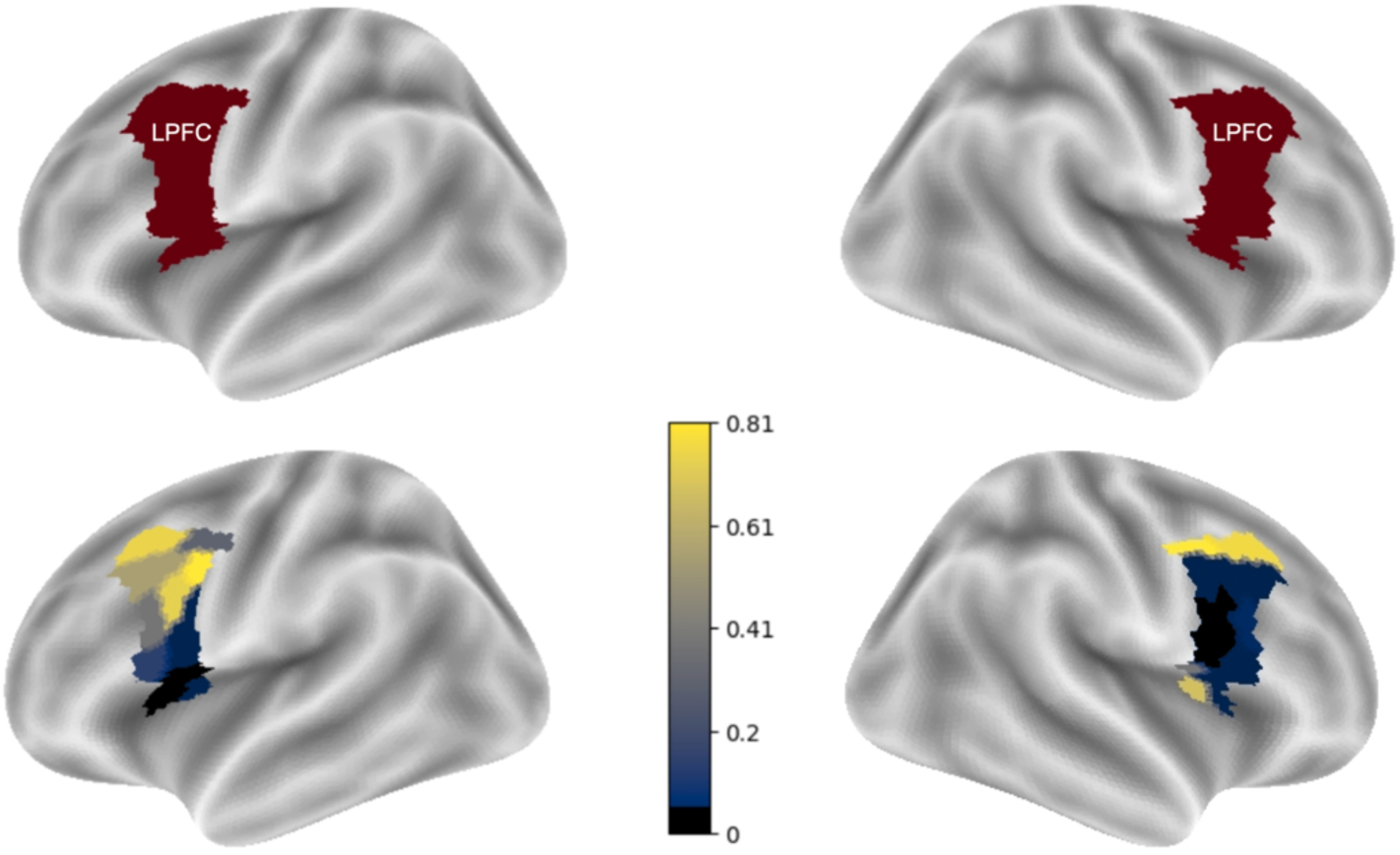
Data-driven control analysis of occlusion-dependent FFA connectivity within lateral prefrontal cortex (LPFC). Top: a LPFC search space in the left and right hemispheres, defined using HCP-MMP parcels associated with lateral prefrontal cortex. Bottom: Parcel-wise significance map of occlusion-dependent FFA connectivity within LPFC. For each LPFC vertex, functional connectivity with FFA was computed using the surface-based 5-mm searchlight distance-correlation procedure. Within each occlusion condition, significant vertices were identified after Bonferroni correction across vertices. For each parcel and each occlusion level, connectivity strength was summarized as the mean correlation across significant vertices; these parcel-level values were then regressed against occlusion severity (5 levels) to obtain a parcel-wise *p* value for the slope term. Parcels surviving Bonferroni correction across LPFC parcels (*p* < .05) are shown in black; all other parcels are colored according to their *p* values.

**Supplementary Figure 7.**
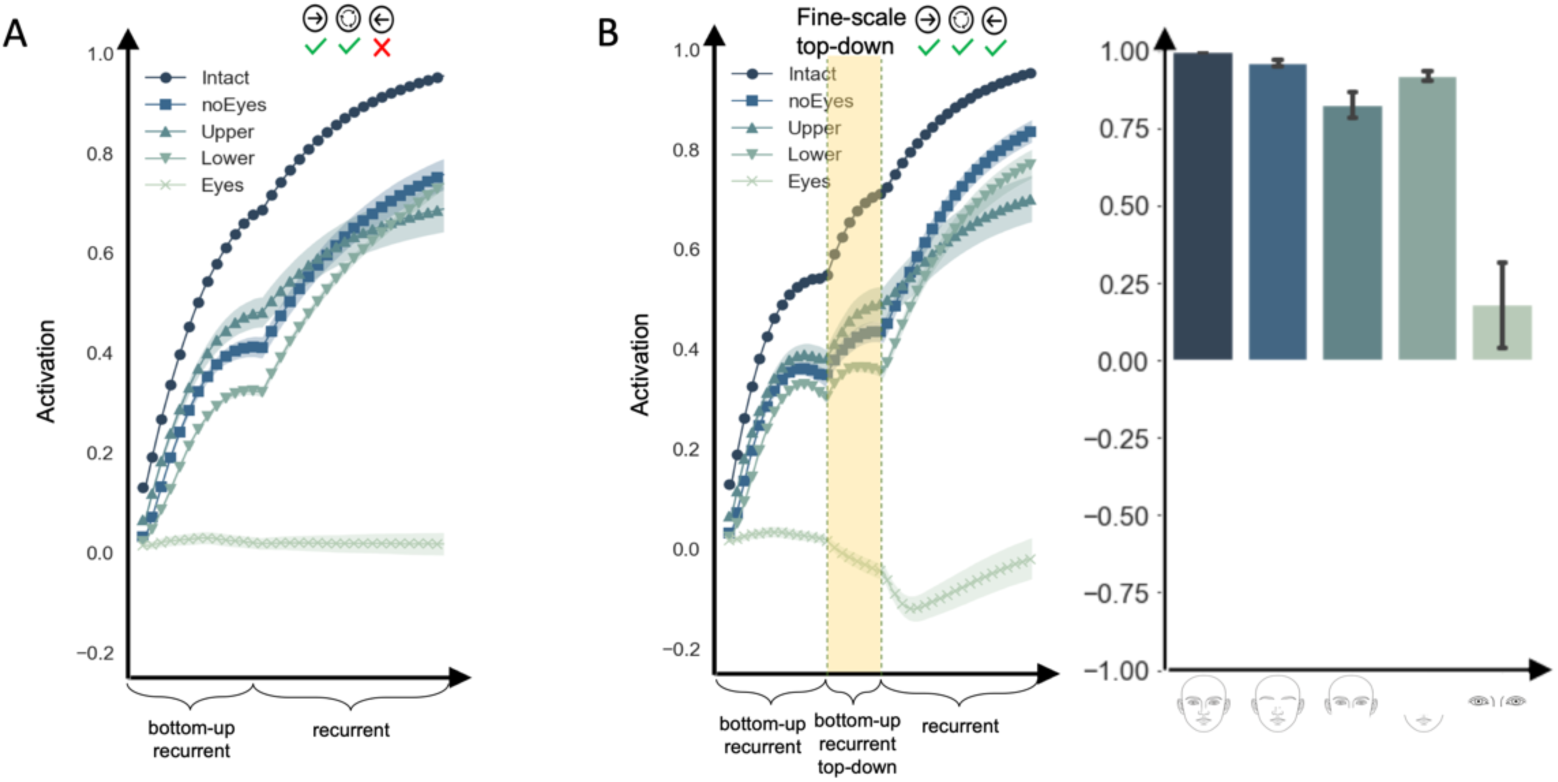
The performance of the control models under different occlusion conditions. (A) Dynamics of the neural state (mean activation in the face region minus mean activation in the object region) in the VTC module without vlPFC feedback in each occlusion condition. Shaded areas indicate standard error (mean ± SEM). (B) Left: dynamics of the neural state in the VTC module with fine-scale feedback. The only difference between this model and the original vlPFC top-down approach lies in training the internal weight connections within the vlPFC using the face-scale representations—specifically face, tool, body, and scene—from the vlPFC. This drives the vlPFC to represent and transmit fine-scale signals to the VTC module. Right: Decoding performance of stable state of the VTC module with or without feedback from vlPFC module.

**Supplementary Figure 8.**
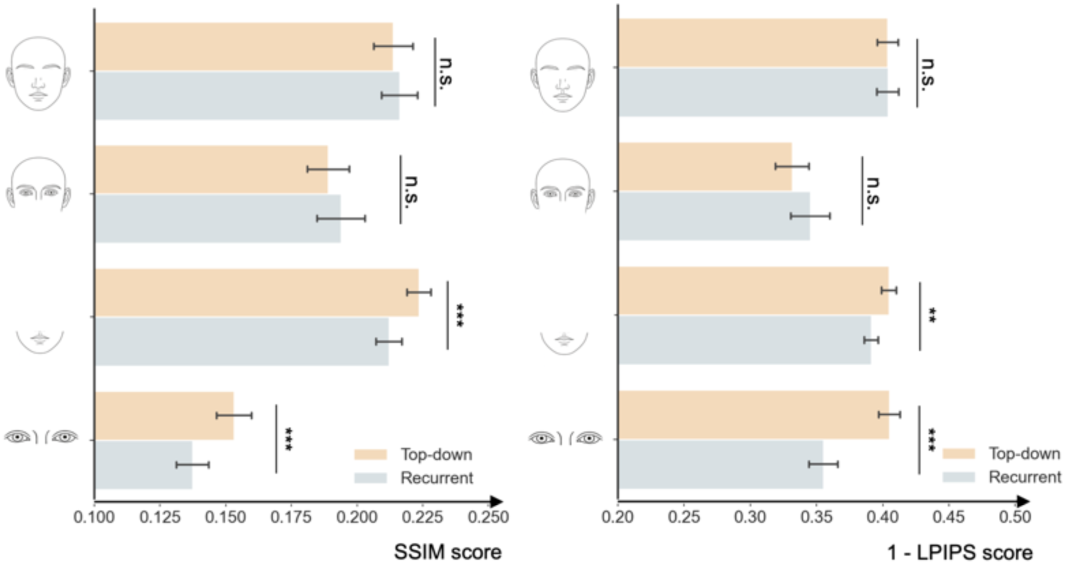
Comparison of image restoration quality under various occlusion conditions with and without top-down information. The left and right panels show the similarity between generated images and actual images measured by the SSIM structural similarity method and the LPIPS semantic similarity method, with and without top-down information. It is found that both SSIM and LPIPS indicate that generated facial images with top-down information exhibit greater similarity to the original images under severe occlusion conditions (SSIM, Lower, *t*(38) = 4.15, *p* < .001; Eyes, *t*(38) = 4.76, *p* < .001; LPIPS, Lower, *t*(38) = 3.03, *p* < .01; Eyes, *t*(38) = 5.69, *p* < .001). **: p < 0.01; ***: p < 0.001

**Supplementary Figure 9.**
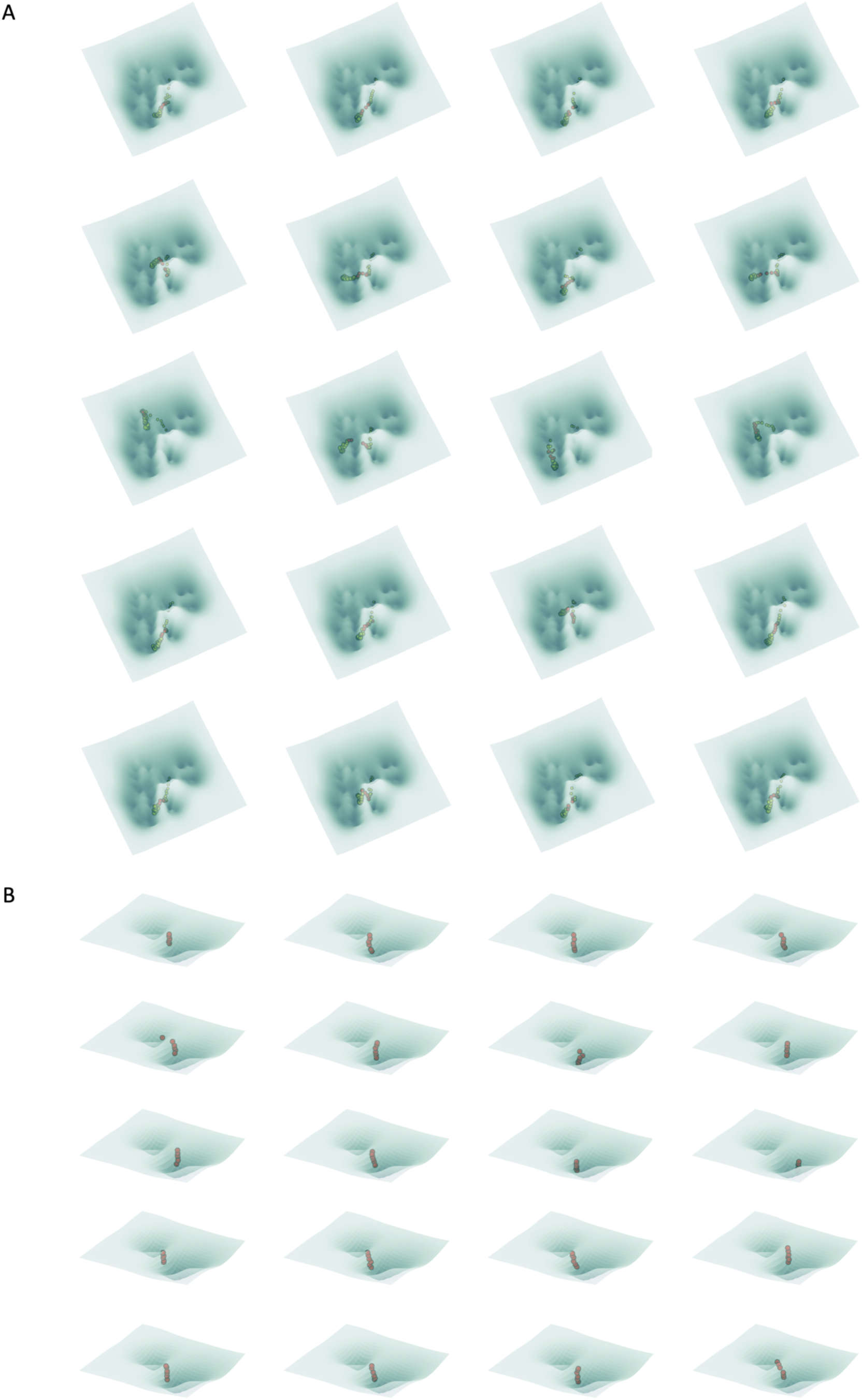
Trajectories of all 20 stimuli under Eyes conditions on the energy-representation manifold. (A) Black dots represent trajectories of VTC module state changes, while red dots indicate behaviors during periods of vlPFC feedback. (B) Red dots denote trajectories of vlPFC module state changes. It can be observed that as the vlPFC state transitions into the animacy attractor, the VTC state gradually emerges from the male-female pseudo-state attractor and moves toward the face attractor.

**Supplementary Figure 10.**
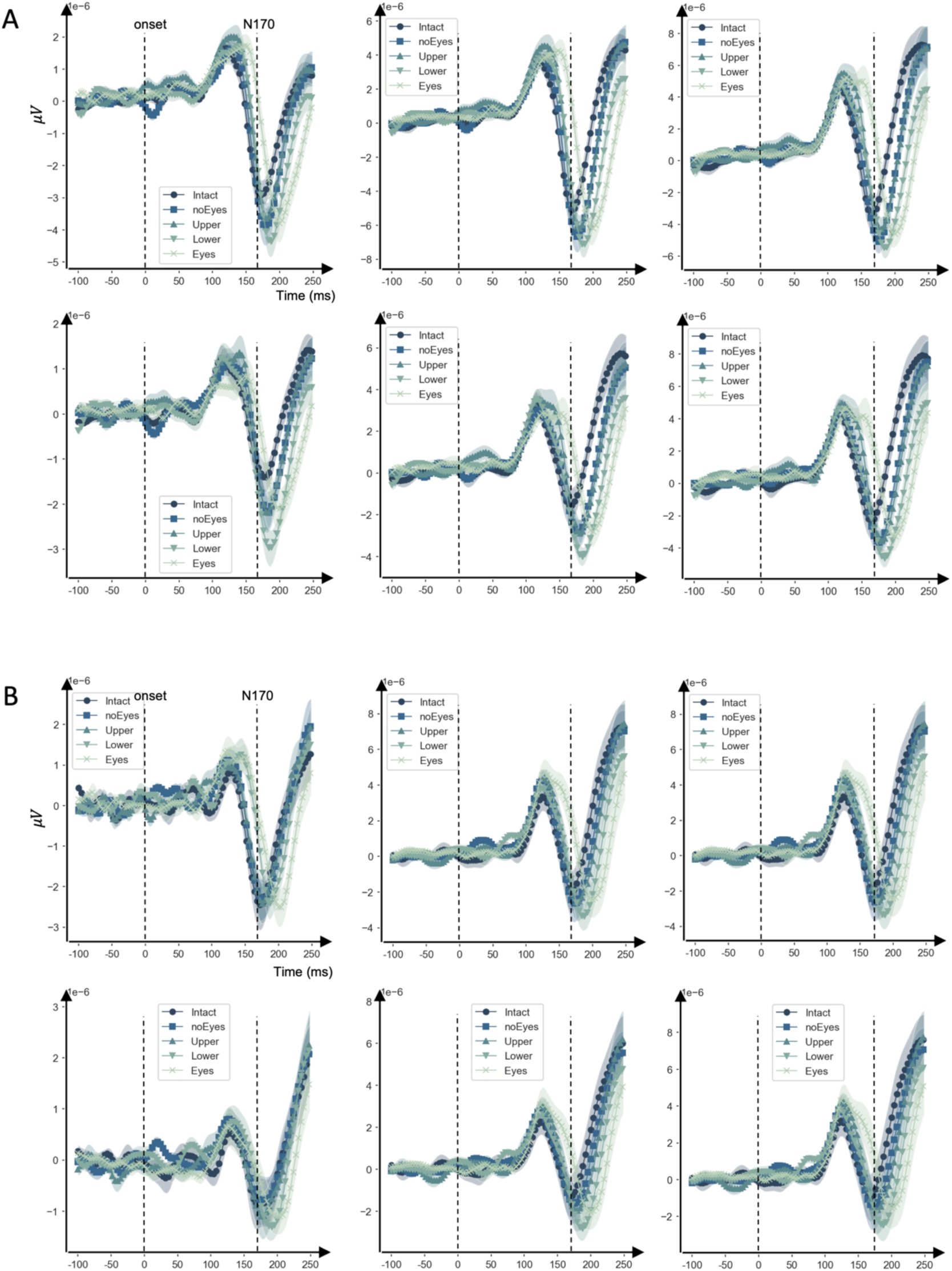
Time series of face channels under different occlusion conditions. After averaging all subjects and trials across different conditions, the time series for each face-selective channel in the left hemisphere (A) and right hemisphere (B) under various occlusion conditions.

**Supplementary Figure 11.**
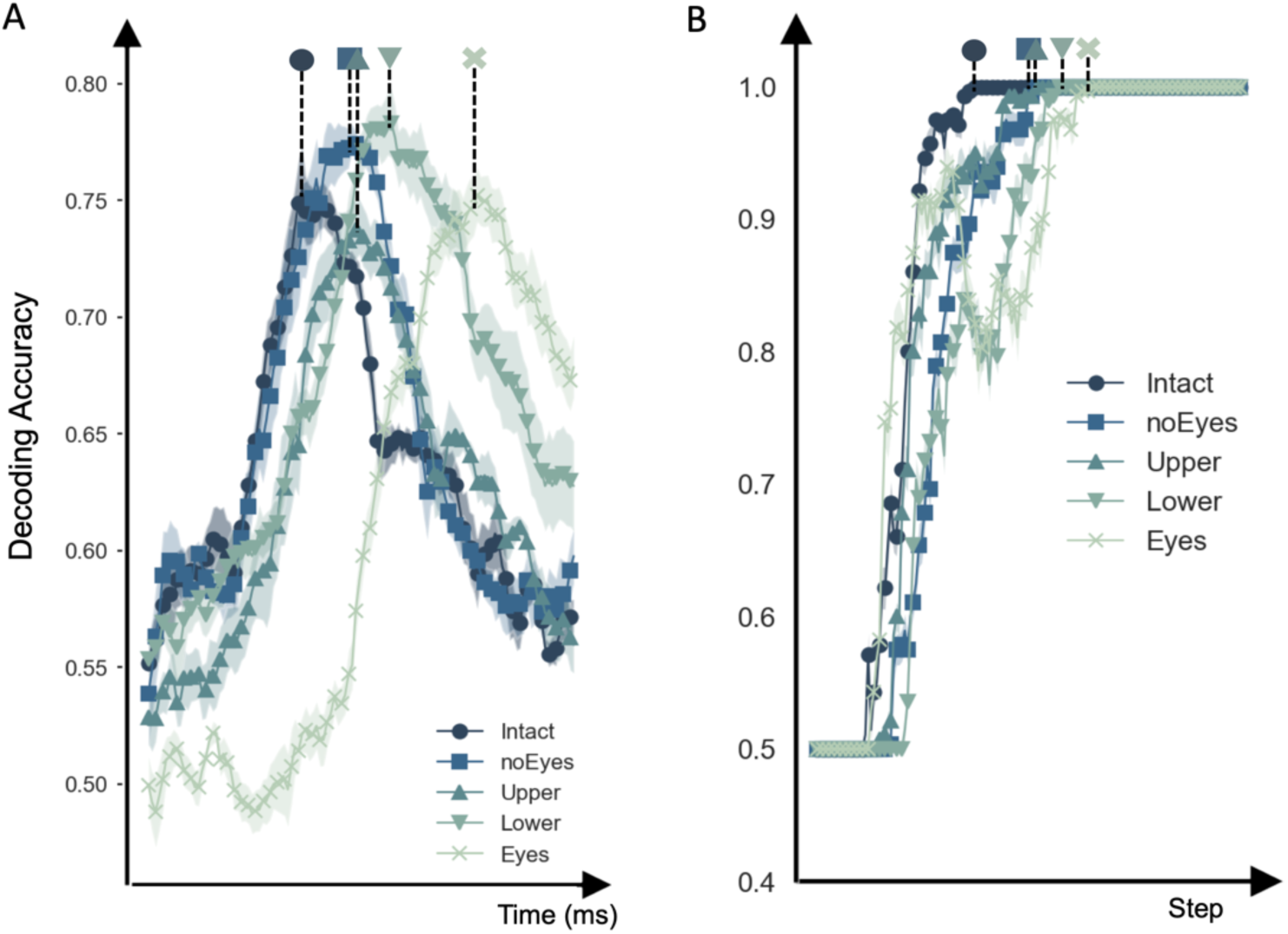
Time series of decoding accuracy under different occlusion conditions. Time series of decoding accuracies of the face-selective channels in EEG (A) and face-selective region in computational model (B) for the five levels of occluded faces. The peak values of each decoding accuracy curve are labeled with different shapes, which can visually show that the EEG and model have consistent hysteresis under different occlusion conditions.

## Reference

1 Felleman, D. J. & Van Essen, D. C. Distributed hierarchical processing in the primate cerebral cortex. Cerebral cortex (New York, NY: 1991) 1, 1–47 (1991).

2 Markov, N. T. et al. Anatomy of hierarchy: feedforward and feedback pathways in macaque visual cortex. Journal of comparative neurology 522, 225–259 (2014).

3 Kubilius, J. et al. Cornet: Modeling the neural mechanisms of core object recognition. BioRxiv, 408385 (2018).

4. Linsley, D., Kim, J., Ashok, A. & Serre, T. Recurrent neural circuits for contour detection. arXiv preprint arXiv:2010.15314 (2020).

5 Dosovitskiy, A. An image is worth 16×16 words: Transformers for image recognition at scale. arXiv preprint arXiv:2010.11929 (2020).

6 Xie, S., Singer, J., Yilmaz, B., Kaiser, D. & Cichy, R. M. Recurrence affects the geometry of visual representations across the ventral visual stream in the human brain. PLoS Biology 23, e3003354 (2025).

7 Briggs, F. Role of feedback connections in central visual processing. Annual review of vision science 6, 313–334 (2020).

8 Velarde, O. M., Makse, H. A. & Parra, L. C. Architecture of the brain’s visual system enhances network stability and performance through layers, delays, and feedback. PLOS Computational Biology 19, e1011078 (2023).

9 Johnson, J. A. & Bullock, D. H. Fragility in AIs Using Artificial Neural Networks. Communications of the ACM 66, 28–31 (2023).

10 Yuille, A. & Kersten, D. Vision as Bayesian inference: analysis by synthesis? Trends in cognitive sciences 10, 301–308 (2006).

11 Poggio, T., Torre, V. & Koch, C. Computational vision and regularization theory. Readings in computer vision, 638–643 (1987).

12 Riesenhuber, M. & Poggio, T. Hierarchical models of object recognition in cortex. Nature neuroscience 2, 1019–1025 (1999).

13 Rolls, E. & Deco, G. Computational neuroscience of vision. (Oxford university press, 2001).

14 Riesenhuber, M. & Poggio, T. Neural mechanisms of object recognition. Current opinion in neurobiology 12, 162–168 (2002).

15 Yamins, D. L. & DiCarlo, J. J. Using goal-driven deep learning models to understand sensory cortex. Nature neuroscience 19, 356–365 (2016).

16 Yamins, D. L. et al. Performance-optimized hierarchical models predict neural responses in higher visual cortex. Proceedings of the national academy of sciences 111, 8619–8624 (2014).

17 Cadieu, C. F. et al. Deep neural networks rival the representation of primate IT cortex for core visual object recognition. PLoS computational biology 10, e1003963 (2014).

18 Khaligh-Razavi, S. M. & Kriegeskorte, N. Deep supervised, but not unsupervised, models may explain IT cortical representation. PLoS Comput Biol 10, e1003915, doi:10.1371/journal.pcbi.1003915 (2014).

19 LeCun, Y. & Bengio, Y. Convolutional networks for images, speech, and time series. The handbook of brain theory and neural networks 3361, 1995 (1995).

20 Tang, H. et al. Recurrent computations for visual pattern completion. Proceedings of the National Academy of Sciences 115, 8835–8840 (2018).

21 Wyatte, D., Curran, T. & O’Reilly, R. The limits of feedforward vision: Recurrent processing promotes robust object recognition when objects are degraded. Journal of cognitive neuroscience 24, 2248–2261 (2012).

22 Johnson, J. S. & Olshausen, B. A. The recognition of partially visible natural objects in the presence and absence of their occluders. Vision research 45, 3262–3276 (2005).

23 Tang, H. et al. Spatiotemporal dynamics underlying object completion in human ventral visual cortex. Neuron 83, 736–748 (2014).

24 Noroozi, J., Rezayat, E. & Dehaqani, M.-R. A. Frontotemporal network contribution to occluded face processing. Proceedings of the National Academy of Sciences 121, e2407457121 (2024).

25 Fyall, A. M., El-Shamayleh, Y., Choi, H., Shea-Brown, E. & Pasupathy, A. Dynamic representation of partially occluded objects in primate prefrontal and visual cortex. Elife 6, e25784 (2017).

26 Choi, H., Pasupathy, A. & Shea-Brown, E. Predictive coding in area V4: dynamic shape discrimination under partial occlusion. Neural computation 30, 1209–1257 (2018).

27 Summerfield, C. et al. Predictive codes for forthcoming perception in the frontal cortex. Science 314, 1311–1314 (2006).

28 Kovács, G., Vogels, R. & Orban, G. A. Selectivity of macaque inferior temporal neurons for partially occluded shapes. Journal of Neuroscience 15, 1984–1997 (1995).

29 Kar, K. & DiCarlo, J. J. Fast recurrent processing via ventrolateral prefrontal cortex is needed by the primate ventral stream for robust core visual object recognition. Neuron 109, 164–176. e165 (2021).

30 Barbas, H. & Mesulam, M. Cortical afferent input to the principals region of the rhesus monkey. Neuroscience 15, 619–637 (1985).

31 Ninomiya, T., Sawamura, H., Inoue, K.-i. & Takada, M. Segregated pathways carrying frontally derived top-down signals to visual areas MT and V4 in macaques. Journal of Neuroscience 32, 6851–6858 (2012).

32 Gerbella, M., Belmalih, A., Borra, E., Rozzi, S. & Luppino, G. Cortical connections of the macaque caudal ventrolateral prefrontal areas 45A and 45B. Cerebral Cortex 20, 141–168 (2010).

33 Miller, E. K. & Cohen, J. D. An integrative theory of prefrontal cortex function. Annual review of neuroscience 24, 167–202 (2001).

34 Bar, M. et al. Top-down facilitation of visual recognition. Proc Natl Acad Sci U S A 103, 449–454, doi:10.1073/pnas.0507062103 (2006).

35 Friston, K. A theory of cortical responses. Philosophical transactions of the Royal Society B: Biological sciences 360, 815–836 (2005).

36 Kok, P. & de Lange, F. P. in An introduction to model-based cognitive neuroscience 221–244 (Springer, 2015).

37 Kanwisher, N., McDermott, J. & Chun, M. M. The fusiform face area: a module in human extrastriate cortex specialized for face perception. Journal of neuroscience 17, 4302–4311 (1997).

38 Tsao, D. Y., Freiwald, W. A., Tootell, R. B. & Livingstone, M. S. A cortical region consisting entirely of face-selective cells. Science 311, 670–674 (2006).

39 Grill-Spector, K., Weiner, K. S., Kay, K. & Gomez, J. The functional neuroanatomy of human face perception. Annual review of vision science 3, 167–196 (2017).

40 Namima, T. & Pasupathy, A. Encoding of partially occluded and occluding objects in primate inferior temporal cortex. Journal of Neuroscience 41, 5652–5666 (2021).

41 Kosai, Y., El-Shamayleh, Y., Fyall, A. M. & Pasupathy, A. The role of visual area V4 in the discrimination of partially occluded shapes. Journal of Neuroscience 34, 8570–8584 (2014).

42 Rajaei, K., Mohsenzadeh, Y., Ebrahimpour, R. & Khaligh-Razavi, S.-M. Beyond core object recognition: Recurrent processes account for object recognition under occlusion. PLoS computational biology 15, e1007001 (2019).

43 Kamps, F. S., Morris, E. J. & Dilks, D. D. A face is more than just the eyes, nose, and mouth: fMRI evidence that face-selective cortex represents external features. Neuroimage 184, 90–100 (2019).

44 Royer, J. et al. Greater reliance on the eye region predicts better face recognition ability. Cognition 181, 12–20 (2018).

45 Moon, J. w. & Yoon, H. S. What is the most important facial part for face recognition? ETRI Journal (2025).

46 Diego-Mas, J. A., Fuentes-Hurtado, F., Naranjo, V. & Alcañiz, M. The influence of each facial feature on how we perceive and interpret human faces. i-Perception 11, 2041669520961123 (2020).

47 Wen, H. et al. Neural Encoding and Decoding with Deep Learning for Dynamic Natural Vision. Cereb Cortex 28, 4136–4160, doi:10.1093/cercor/bhx268 (2018).

48 Xu, S., Zhang, Y., Zhen, Z. & Liu, J. The face module emerged in a deep convolutional neural network selectively deprived of face experience. Frontiers in computational neuroscience 15, 626259 (2021).

49 Huang, T., Zhen, Z. & Liu, J. Semantic relatedness emerges in deep convolutional neural networks designed for object recognition. Frontiers in computational neuroscience 15, 625804 (2021).

50 Liu, X., Zhen, Z. & Liu, J. Hierarchical Sparse Coding of Objects in Deep Convolutional Neural Networks. Front Comput Neurosci 14, 578158, doi:10.3389/fncom.2020.578158 (2020).

51 Selvaraju, R. R. et al. in Proceedings of the IEEE international conference on computer vision. 618–626.

52 Krizhevsky, A., Sutskever, I. & Hinton, G. E. Imagenet classification with deep convolutional neural networks. Advances in neural information processing systems 25, 1097–1105 (2012).

53 Naseer, M. M. et al. Intriguing properties of vision transformers. Advances in Neural Information Processing Systems 34, 23296–23308 (2021).

54 Gilbert, C. D. & Li, W. Top-down influences on visual processing. Nature Reviews Neuroscience 14, 350–363 (2013).

55 Glasser, M. F. et al. A multi-modal parcellation of human cerebral cortex. Nature 536, 171–178 (2016).

56 Chung, S., Lee, D. D. & Sompolinsky, H. Classification and geometry of general perceptual manifolds. Physical Review X 8, 031003 (2018).

57 Zhang, Y., Liu, J. & Liu, J. Structured interconnectivity optimizes neural geometry for balancing specificity and generalization in object recognition. Communications Biology (2025).

58 Bugatus, L., Weiner, K. S. & Grill-Spector, K. Task alters category representations in prefrontal but not high-level visual cortex. Neuroimage 155, 437–449 (2017).

59 McKee, J. L., Riesenhuber, M., Miller, E. K. & Freedman, D. J. Task dependence of visual and category representations in prefrontal and inferior temporal cortices. Journal of Neuroscience 34, 16065–16075 (2014).

60 Friedman, N. P. & Robbins, T. W. The role of prefrontal cortex in cognitive control and executive function. Neuropsychopharmacology 47, 72–89 (2022).

61 Konkle, T. & Oliva, A. A real-world size organization of object responses in occipitotemporal cortex. Neuron 74, 1114–1124 (2012).

62 Long, B., Yu, C.-P. & Konkle, T. Mid-level visual features underlie the high-level categorical organization of the ventral stream. Proceedings of the National Academy of Sciences 115, E9015–E9024 (2018).

63 Konkle, T. & Caramazza, A. Tripartite organization of the ventral stream by animacy and object size. J Neurosci 33, 10235–10242, doi:10.1523/JNEUROSCI.0983-13.2013 (2013).

64 Haxby, J. V. et al. Distributed and overlapping representations of faces and objects in ventral temporal cortex. Science 293, 2425–2430 (2001).

65 Schwiedrzik, C. M. & Freiwald, W. A. High-level prediction signals in a low-level area of the macaque face-processing hierarchy. Neuron 96, 89–97. e84 (2017).

66 Lee, T. S. & Mumford, D. Hierarchical Bayesian inference in the visual cortex. Journal of the Optical Society of America A 20, 1434–1448 (2003).

67. Mirza, M. & Osindero, S. Conditional generative adversarial nets. arXiv preprint arXiv:1411.1784 (2014).

68 Ahn, S., Adeli, H. & Zelinsky, G. J. The attentive reconstruction of objects facilitates robust object recognition. PLOS Computational Biology 20, e1012159 (2024).

69 Muckli, L. et al. Contextual feedback to superficial layers of V1. Current Biology 25, 2690–2695 (2015).

70 Liu, J., Higuchi, M., Marantz, A. & Kanwisher, N. The selectivity of the occipitotemporal M170 for faces. Neuroreport 11, 337–341 (2000).

71 Liu, J., Harris, A. & Kanwisher, N. in Social neuroscience 75-85 (Psychology Press, 2013).

72 Millidge, B., Seth, A. & Buckley, C. L. Predictive coding: a theoretical and experimental review. arXiv preprint arXiv:2107.12979 (2021).

73 Mumford, D. On the computational architecture of the neocortex: II The role of cortico-cortical loops. Biological cybernetics 66, 241–251 (1992).

74 Bello, I., Zoph, B., Vaswani, A., Shlens, J. & Le, Q. V. in Proceedings of the IEEE/CVF international conference on computer vision. 3286–3295.

75 Elharrouss, O., Damseh, R., Belkacem, A. N., Badidi, E. & Lakas, A. Transformer-based image and video inpainting: current challenges and future directions. Artificial Intelligence Review 58, 124 (2025).

76 Huang, G. B., Mattar, M., Berg, T. & Learned-Miller, E. in Workshop on faces in’Real-Life’Images: detection, alignment, and recognition.

77 Griffin, G., Holub, A. & Perona, P. Caltech-256 object category dataset. (2007).

78 Van Essen, D. C. et al. The WU-Minn human connectome project: an overview. Neuroimage 80, 62–79 (2013).

79. Simonyan, K. & Zisserman, A. Very deep convolutional networks for large-scale image recognition. arXiv preprint arXiv:1409.1556 (2014).

80. He, K., Zhang, X., Ren, S. & Sun, J. in Proceedings of the IEEE conference on computer vision and pattern recognition. 770–778.

81 Szegedy, C., Vanhoucke, V., Ioffe, S., Shlens, J. & Wojna, Z. in Proceedings of the IEEE conference on computer vision and pattern recognition. 2818–2826.

82 Esteban, O. et al. fMRIPrep: a robust preprocessing pipeline for functional MRI. Nature methods 16, 111–116 (2019).

83 Worsley, K. J. et al. A unified statistical approach for determining significant signals in images of cerebral activation. Human brain mapping 4, 58–73 (1996).

84 Zhen, Z. et al. Quantifying interindividual variability and asymmetry of face-selective regions: a probabilistic functional atlas. Neuroimage 113, 13–25 (2015).

85 Grill-Spector, K. & Weiner, K. S. The functional architecture of the ventral temporal cortex and its role in categorization. Nature Reviews Neuroscience 15, 536–548 (2014).

86 Zhang, Y., Zhou, K., Bao, P. & Liu, J. A biologically inspired computational model of human ventral temporal cortex. Neural Networks, 106437 (2024).

87 Weiner, K. S. et al. The mid-fusiform sulcus: a landmark identifying both cytoarchitectonic and functional divisions of human ventral temporal cortex. Neuroimage 84, 453–465 (2014).

88 Russakovsky, O. et al. Imagenet large scale visual recognition challenge. International journal of computer vision 115, 211–252 (2015).

89 Sherman, S. M. & Guillery, R. The role of the thalamus in the flow of information to the cortex. Philosophical Transactions of the Royal Society of London. Series B: Biological Sciences 357, 1695–1708 (2002).

90 Sillito, A. M. & Jones, H. E. Corticothalamic interactions in the transfer of visual information. Philosophical Transactions of the Royal Society of London. Series B: Biological Sciences 357, 1739–1752 (2002).

91 McInnes, L., Healy, J. & Melville, J. Umap: Uniform manifold approximation and projection for dimension reduction. arXiv preprint arXiv:1802.03426 (2018).

92 Hopfield, J. J. Neural networks and physical systems with emergent collective computational abilities. Proceedings of the national academy of sciences 79, 2554–2558 (1982).

93 Mao, X. et al. in Proceedings of the IEEE international conference on computer vision. 2794–2802.

94. Johnson, J., Alahi, A. & Fei-Fei, L. in European conference on computer vision. 694–711 (Springer).

95 Wang, Z., Bovik, A. C., Sheikh, H. R. & Simoncelli, E. P. Image quality assessment: from error visibility to structural similarity. IEEE transactions on image processing 13, 600–612 (2004).

96. Zhang, R., Isola, P., Efros, A. A., Shechtman, E. & Wang, O. in Proceedings of the IEEE conference on computer vision and pattern recognition. 586–595.

